# Functional Annotation of Chemical Libraries across Diverse Biological Processes

**DOI:** 10.1101/112557

**Authors:** Jeff S. Piotrowski, Sheena C. Li, Raamesh Deshpande, Scott W. Simpkins, Justin Nelson, Yoko Yashiroda, Jacqueline M. Barber, Hamid Safizadeh, Erin Wilson, Hiroki Okada, Abraham A. Gebre, Karen Kubo, Nikko P. Torres, Marissa A. LeBlanc, Kerry Andrusiak, Reika Okamoto, Mami Yoshimura, Eva DeRango-Adem, Jolanda van Leeuwen, Katsuhiko Shirahige, Anastasia Baryshnikova, Grant W. Brown, Hiroyuki Hirano, Michael Costanzo, Brenda Andrews, Yoshikazu Ohya, Hiroyuki Osada, Minoru Yoshida, Chad L. Myers, Charles Boone

## Abstract

Chemical-genetic approaches offer the potential for unbiased functional annotation of chemical libraries. Mutations can alter the response of cells to a compound, revealing chemical-genetic interactions that can elucidate a compound’s mode of action. We developed a highly parallel and unbiased yeast chemical-genetic screening system involving three key components. First, in a drug-sensitive genetic background, we constructed an optimized, diagnostic mutant collection that is predictive all major yeast biological processes. Second, we implemented a multiplexed (768-plex) barcode sequencing protocol, enabling assembly of thousands of chemical-genetic profiles. Finally, based on comparison of the chemical-genetic profiles with a compendium of genome-wide genetic interaction profiles, we predicted compound functionality. Applying this high-throughput approach, we screened 7 different compound libraries and annotated their functional diversity. We further validated biological process predictions, prioritized a diverse set of compounds, and identified compounds that appear to have dual modes of action.

## INTRODUCTION

Discovery and development of novel compound libraries has outpaced the functional characterization of these compounds, leading to a growing knowledge gap^1,2^. Chemical probes that target specific cellular functions are valuable entities because they can provide insight into fundamental cellular functions and represent putative leads for new drug development. Despite a massive wealth of whole-genome sequence data that has identified hundreds of potential new druggable targets, in both humans and pathogens, we lack the chemical probes to take advantage of these insights^3^. Therefore, a major demand exists for large-scale functional annotation of bioactive compounds.

Whole-cell screening approaches are advantageous because they identify bioavailable molecules and provide readouts based on general bioactivity^4^, a particular phenotypic response^5^, or a specific reporter system^6^ while maintaining biological context. Chemical-genetics expands traditional whole cell screening because it has the potential to monitor all cellular pathways in an unbiased manner^7,8^. A typical chemical-genetic screen involves testing a collection of mutant strains with defined genetic perturbations for fitness defects or advantages when grown in the presence of a specific compound^9–11^. Quantifying the relative fitness of a collection of mutant strains in response to compound treatment generates a chemical-genetic interaction profile, which provides diagnostic functional information about a compound’s general mode-of-action^10,12^.

*Saccharomyces cerevisiae* represents a powerful eukaryotic model system for chemical-genetic analysis, due to its facile genetics and availability of functional genomic reagents and tools. For example, genome-wide gene deletion analysis^13^, identified ~1000 essential genes and enabled the generation of a set of ~5000 viable haploid deletion mutants. The essential genes can be exploited for chemical genetic studies as heterozygous diploid mutants, whereas the nonessential genes can be studied as viable haploid deletion mutants, such that each mutant is examined for hypersensitivity or resistance to a compound^10, 11^. Each strain is uniquely barcoded allowing the responses of hundreds of pooled mutants to be measured simultaneously to generate a chemical-genetic interaction profile^9,10^.

A comprehensive genetic interaction network, in which the majority of all possible double mutants are scored for genetic interactions quantitatively, has been mapped for yeast^14^. A genetic interaction occurs when mutations in two or more genes combine to generate an unexpected phenotype. Given the single mutant phenotypes, a negative genetic interaction occurs when two mutations combine to produce a double mutant fitness defect that is more severe than expected, whereas a positive genetic interaction reflects a double mutant fitness defect that is less severe than expected. The set of negative and positive genetic interactions for a particular query gene represents a genetic interaction profile, which provides a quantitative description of gene function. A global network of genetic interaction profile similarities groups genes with similar roles into dense gene clusters that represent major biological processes and thus highlights the functional organization of a cell^14, 15^. Importantly, a global compendium of genetic interaction profiles can be used to functionally interpret chemical-genetic interaction profiles^12,15^. If a bioactive compound inhibits a specific target protein, then loss-of-function mutations in the corresponding target gene should mimic the bioactivity of the compound^12,15^. Moreover, the genetic interaction profile of the target gene should resemble the chemical-genetic interaction profile of the inhibitory compound that modulates the target pathway^12,15^. For example, the genetic interaction profile associated with a partial loss-of-function mutation in *ERG11,* which encodes the target of fluconazole, closely resembles the chemical-genetic interaction profile of fluconazole^12^. Thus, the global genetic interaction network provides a general key for interpreting the target pathways of bioactive compounds, enabling compounds to be annotated to specific biological processes and possibly specific pathways.

We developed a high-throughput chemical-genetic screening platform to functionally annotate large compound collections in a rapid and systematic manner. To do so, we constructed a diagnostic set of viable yeast gene deletion mutants, each carrying a unique DNA barcode identifier, which span all major biological processes, within a drug-sensitized, genetic background. We also developed a highly multiplexed (768-plex) barcode sequencing protocol, allowing us to decipher rich chemical-genetic profiles for hundreds of compounds simultaneously. Finally, we assembled a computational platform for functionally annotating compounds to specific biological processes and pathways. Ultimately, we applied this chemical-genetic pipeline to annotate seven diverse libraries containing 13,524 compounds in an unbiased and systematic manner.

## RESULTS

To design a pipeline for high-throughput chemical-genetic profiling and functional annotation of chemical libraries (**Fig. 1a**), we first selected an optimal set of diagnostic genes and constructed a mutant strain collection in which each diagnostic gene was individually deleted in a drug-hypersensitive genetic background. Second, we developed a highly multiplexed barcode sequencing^16^ system for chemical-genetic profiling with optimized signal detection. Third, we implemented computational approaches to integrate chemical-genetic profiles with the global yeast genetic interaction network to predict biological processes targeted by specific compounds. Finally, we assembled a database of chemical structures, chemical-genetic profiles, and functional predictions for each library investigated in this study.

**Figure 1.**
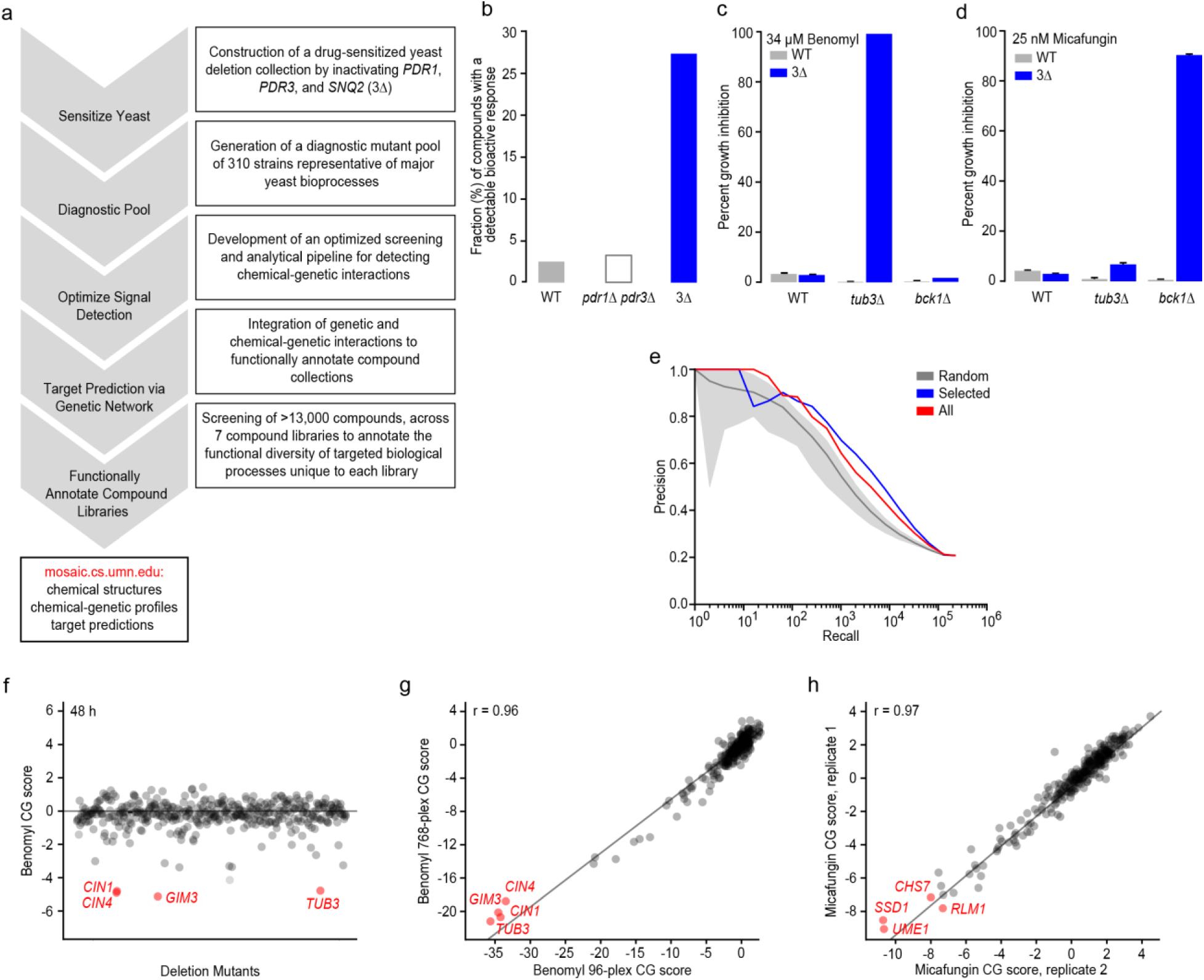
Miniaturizing chemical-genetic profiling. **(a)** A high-throughput chemical-genetics platform for functional annotation of compound libraries. **(b)** The fraction (%) of compounds showing a bioactive response based on detection of a halo of growth inhibition surrounding a compound spotted on a lawn of WT strain, a *pdr1∆ pdr3∆* double mutant, or a *pdr1∆ pdr3∆ snq2∆* triple mutant strain (3Δ). **(c)** Comparison of WT vs. 3Δ strains for detecting a benomyl*-TUB3* chemical-genetic interaction (n=3, mean ± S.E.). (**d**) Comparison of WT vs. 3Δ strains for detecting a micafungin-*BCK1* chemical-genetic interaction (n=3, mean ± S.E.). **(e)** Plots of precision [True positives / (True positives + False positives)] versus recall (total number of true positives) to evaluate gene function predictions based on genetic interaction profile similarities derived from the entire non-essential deletion mutant collection (red), the diagnostic strain collection (blue), and a random selection of deletion strains the same size as the diagnostic collection (grey). True positives were defined as those gene pairs where both genes are annotated to the same GO gold standard set of terms^72^. **(f)** Detection of chemical-genetic interactions (red) following 48 h growth in the presence of benomyl. **(g)** Correlation of average benomyl chemical-genetic interaction profiles (n=3, technical replicates) derived from multiplexing 96 vs. 768 chemical genetic screens in a single sequencing lane. Benomyl-specific chemical-genetic interactions are shown in red. **(h)** Correlation of micafungin chemical-genetic interaction profiles derived from two independent biological replicates. Specific micafungin chemical-genetic interactions are shown in red.

### Developing a diagnostic gene set for chemical-genetic profiling

To increase the potential for detecting bioactive compounds, we constructed a drug-sensitized yeast genetic background by combining deletions of *PDR1* and *PDR3*, both of which encode transcription factors known to regulate the yeast pleiotropic drug response^17,18^, with a deletion of *SNQ2,* which encodes a multidrug transporter (**Supplementary Results**, **Supplementary Fig. 1**). We tested growth of the resultant *pdr1∆ pdr3∆ snq2∆* (3Δ) drug-sensitized strain in the presence of 440 different control compounds (see **Methods**, **Supplementary Table 1**) and observed a ~5-fold increase in the number of compounds that inhibited growth of the drug-sensitized strain compared to wild-type cells via a halo assay, indicating that these deletion mutations sensitized yeast to diverse classes of compounds (**Fig. 1b**)^19^. When considering the complete set of 13,524 compounds tested in this study, the average “hit rate”, corresponding to the fraction of bioactive compounds within a collection that causes at least 20% growth inhibition in the drug-sensitized strain in liquid medium, was ~35% across all compounds tested, which is ~5X greater than the hit rate found using the equivalent wild-type strain background in previous studies^11^ (**Supplementary Table 1**). Specific chemical-genetic interactions were also detected more readily in the drug-sensitized background. For example, at a concentration of 34.4 μΜ, the microtubule-binding compound benomyl showed a specific chemical-genetic interaction with *TUB3,* which encodes α-tubulin, only in our drug-sensitized background (**Fig. 1c**). Similarly, we analyzed the response to a cell wall glucan synthase inhibitor, micafungin, at 25 nM, and we detected a specific chemical-genetic interaction with *BCK1,* which encodes a component of the PKC cell wall integrity-signaling pathway (**Fig. 1d**). In both cases, only the known sensitive mutant showed an exaggerated chemical-genetic interaction, suggesting that, like wild-type cells, the drug-sensitive background identifies functionally relevant signals (**Fig. 1c-d**).

Because genes within the same pathway and the same biological process tend to share similar genetic interaction profiles^12,15^, only a subset of genes are required to capture functionally informative genetic interaction signatures for a given gene. Leveraging this property, we developed a computational approach for optimal selection of mutants for chemical-genetic screens, identifying a set of 157 functionally diagnostic strains (**Fig. 1e)** (see **Methods**). Independently, we also manually selected 236 strains mutated for genes that span major yeast biological processes that belong to highly-connected clusters in the global genetic interaction profile similarity network^15^, 83 of which overlapped with the computationally selected set. Thus, the final diagnostic pool consisted of 310 deletion mutant strains (~6% of all nonessential genes) that spans a similar functional space as the entire non-essential deletion mutant collection (**Supplementary Fig. 2, Supplementary Table 2**). While members of our diagnostic subset are not distributed proportionally across the 17 major bioprocesses, these were selected not only for bioprocess representation, but also their predictive power (see **Methods**). Even though we are using a subset of strains, this diagnostic collection has been optimized for gene similarity-based target prediction across the entire set of genetic interaction query strains (**Fig. 1e**).

Furthermore, we compared the individual fitness of each sensitized deletion strain to the original deletion collection (**Supplementary Table 2**), and used this fitness score to select pool members with near-equivalent fitness. We observed that ~20% of the mutants in diagnostic pool version 2.0 could not be scored by our standard SGA scoring method because of irregularities in colony shape in the *pdr1∆ pdr3∆ snq2∆* genetic background (**Supplementary Table 2**), and we verified that these mutants had appropriate fitness values based on barcode representation after pooled liquid growth. Reducing the complexity of the ~5000 viable yeast deletion mutant collection to a smaller diagnostic set allowed us to maximize the dynamic range for detecting chemical-genetic interactions in a micro-culture and increased the degree of multiplexing for our barcode sequencing read-out.

### Optimizing signal detection and high-throughput screening

Detecting drug-gene interactions requires a clear separation of sensitive/resistant mutants relative to the unaffected mutants in the pooled assay. To optimize signal detection, we tested the effects of three factors on detection of drug-gene interactions using the well-characterized compounds benomyl and micafungin. These included inoculum size, incubation time, and the number of PCR cycles used for barcode DNA amplification (see **Methods**). Incubation time had the most pronounced effect on the signal to noise ratio of the chemical-genetic profiles, with the optimal outcome observed after 48 h incubation (**Fig. 1f, Supplementary Fig. 3a**). For example, gene deletion mutants defective in microtubule functions, including *CIN1, CIN4, GIM3* and *TUB3,* were depleted efficiently from the culture after 48 h growth in the presence of benomyl. The assay was relatively robust to inoculum density and number of PCR amplification cycles (**Supplementary Fig. 3a**). Ultimately, the screening conditions we selected included 200 μL micro-cultures, 48 h growth, at an inoculum of 250 cells/strain and 30 PCR cycles for barcode amplification. These parameters resulted in high correlation between biological replicates (**Supplementary Fig. 3b**).

Multiplexing of chemical-genetic samples is critical for screening large chemical libraries composed of thousands of compounds. Employing a custom-designed set of 768 multiplex primers, each containing a unique 10bp multiplex tag (**Supplementary Table 3,** see **Methods**), we found that combining the barcode DNA samples from 768 different chemical-genetic experiments produced profiles of similar quality to profiles for the same set of compounds generated at 96-plex (**Fig. 1g**). Thus, we adopted a screening strategy of 768 samples per Illumina HiSeq sequencing lane, or 6144 samples per flow cell. Under these conditions, biological replicates (independently grown cultures of the same strain pool) from different sequencing lanes exhibited highly reproducible chemical-genetic profiles (**Fig. 1h**). In pilot experiments, we sequenced barcodes using two separate reads, one for the multiplex tag and another for the deletion barcode; this methodology was thought to improve sequencing accuracy because it reduces the read length^20^. However, we achieved a more uniform distribution of sequence counts across conditions and barcoded mutants by using a single sequencing reaction designed to read through the entire PCR amplicon (**Supplementary Fig. 3c**).

### Chemical-genetic profiling of diverse compound libraries

Applying our optimized pipeline, we generated chemical-genetic interaction profiles for 13,524 compounds by screening seven diverse compound collections: the RIKEN Natural Product Depository (NPDepo), which is composed largely of purified natural products or natural product derivatives, four collections from the National Cancer Institute’s Open Chemical Repository (natural products: NCI-NP, approved oncology drugs: NCI-ONC, structural and mechanistic diversity sets: NCI-STRUCT-DIV and NCI-MECH-DIV, respectively), a library of compounds from the National Institutes of Health Small Molecule Repository with a history of use in human clinical trials (NIH Clinical Collection or NIHCC), and the Glaxo-Smith-Klein kinase inhibitor collection (GSK-KI). A complete description of these collections, all compounds screened, their structures, basic physical properties, and chemical-genetic data is provided (**Supplementary Table 4** and **Supplementary Information**).

Chemical-genetic interactions were identified and scored by comparing the individual mutant barcode read counts to those from a set of solvent control conditions. A negative chemical genetic (CG) interaction score represents hypersensitivity to a compound whereas a positive CG score represents resistance (see **Methods**). At a relatively strict CG score threshold of +/-2.5 (z-score for enrichment/depletion in the presence of the compound relative to DMSO control), we observed positive chemical-genetic interactions between 0.5% of all compound-deletion mutant pairs, and negative chemical-genetic interactions between 1.1% of all compound-deletion mutant pairs. The set of highly bioactive compounds, which inhibited growth of the pooled collection by more than 20% (~4700 compounds), exhibited a substantially higher frequency of chemical-genetic interactions, with 1.3% and 2.3% of compound-mutant pairs for positive and negative interactions, respectively. Each deletion mutant displayed, on average, ~46 positive interactions and ~79 negative interactions across the entire collection of screened compounds. The number of chemical-genetic interactions for each strain (CG score ≥ 2.5 or ≤ −2.5) across all screened compounds is presented in **Supplementary Table 5**. Importantly, compounds screened both in our study, using the diagnostic set and in previous studies using the entire nonessential deletion mutant collection showed positive correlations, despite differences in strain backgrounds and methods used to measure mutant strain abundance (microarray vs. sequencing) (**Supplementary Table 6**).^11,21^

Hierarchical clustering analysis^10,12^ provides a visual representation of the diversity of the resultant chemical-genetic profiles. We focused on the most responsive subset of 173 gene deletion mutants, whose chemical-genetic profiles consisted of at least three extreme negative interactions (CG score ≤ −5), and 1380 compounds, which were derived from all seven collections (**Fig. 2**, See **methods**). The clustered matrix highlighted chemical-genetic interactions involving sets of functionally related genes participating in different biological processes, including DNA replication & repair (**i**), mitosis and chromosome segregation (**ii**), glycosylation, protein folding/targeting, cell wall biogenesis (**iii**), transcription and chromatin organization (**iv**), vesicle traffic (**v**), cell polarity and morphogenesis (**vi**), and other biological functions (**Fig. 2**). For example, a cluster of compounds, including benomyl and the tubulin-binding compound nocodazole, showed specific chemical-genetic interactions with *TUB3* and *CIN1,* suggesting these compounds may target microtubule function or, more generally, target pathways with roles in mitosis and chromosome segregation. Indeed, this cluster includes a previously uncharacterized compound from the RIKEN NPDepo collection, NPD2784, which we found strongly inhibits polymerization of mammalian tubulin *in vitro* (**Supplementary Fig. 4**).

**Figure 2.**
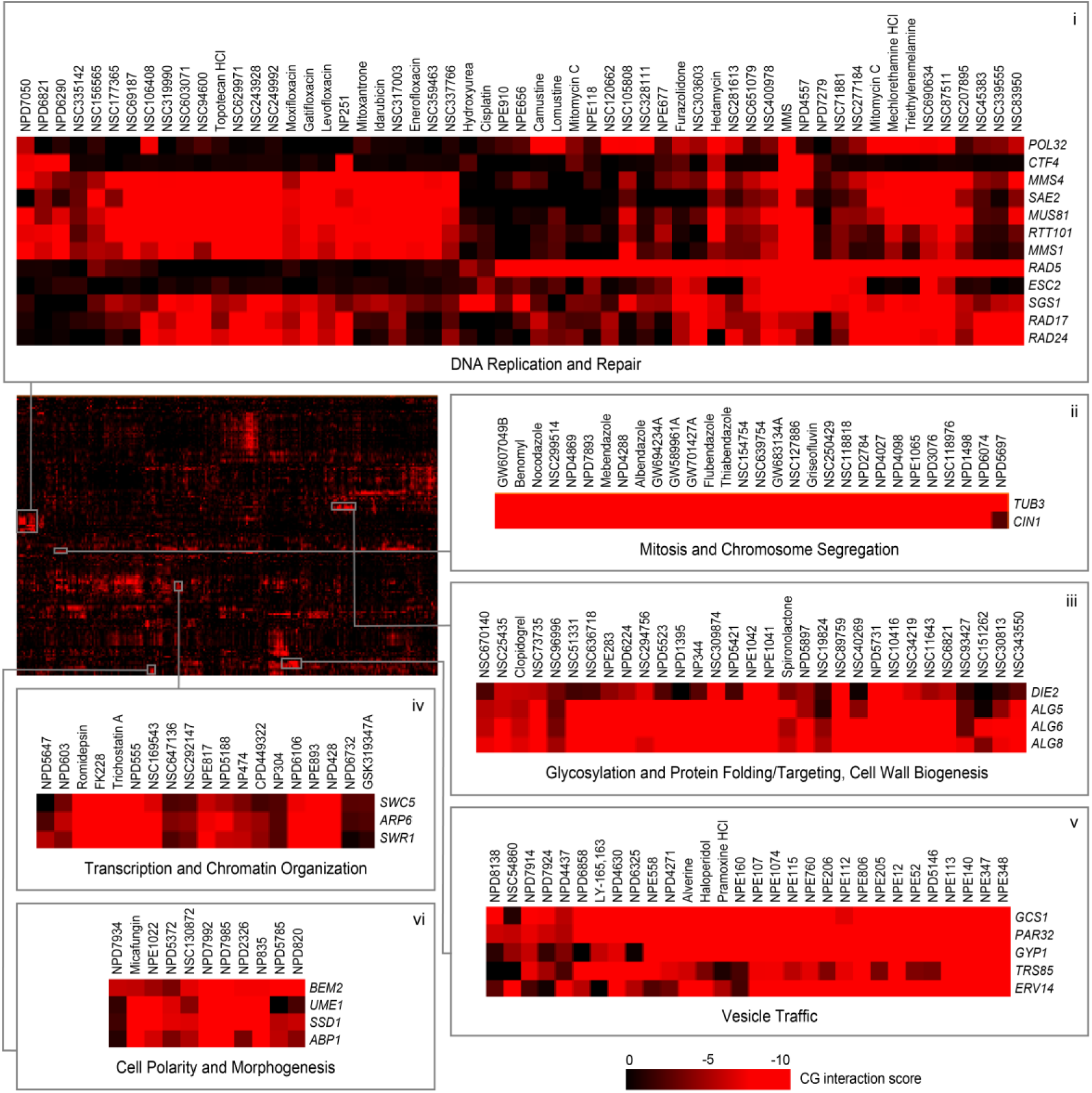
Two-dimensional hierarchical clustering of chemical-genetic interactions. Mean negative chemical-genetic interactions are represented in red (n=3, technical replicates). Rows, 173 deletion mutant strains; columns, 1380 bioactive compounds from the high confidence set (HCS). Sections are expanded to allow detailed visualization of compounds targeting processes related to DNA replication & repair (**i**), mitosis and chromosome segregation (**ii**), glycosylation, protein folding/targeting, and cell wall biogenesis (**iii**), transcription and chromatin organization (**iv**), vesicle traffic (**v**), cell polarity and morphogenesis (**vi**).

### Integrating genetic and chemical-genetic profiles

The chemical-genetic interaction profile of a compound that targets a specific biological process should overlap the genetic interaction profiles of genes that function as part of that biological process^12,15^. To identify biological processes targeted by compounds, we compared the chemical-genetic profile of each compound to our comprehensive set of genetic interaction profiles (**Supplementary Fig. 5**), allowing us to score each compound-gene pair for profile similarity (see **Methods**). This analysis generated a set of gene-level similarity scores identifying a set of potential target genes for each compound. Although prediction of the precise gene target requires deeper experimental analysis, our approach readily predicted the biological process targeted by a particular compound based on Gene Ontology (GO) annotations shared among the target gene set (see **Methods**). To focus on high-confidence predictions, we estimated false discovery rates (FDR) for biological process-level predictions based on both resampled and DMSO control profiles and applied specific FDR thresholds (RIKEN NPDepo screen: FDR ≤ 25%; NCI/NIH/GSK screen: FDR ≤ 27%, see **Methods**). This analysis yielded 1522 high-confidence compound profiles that we refer to as our high confidence set (HCS) (**Supplementary Table 7**). We found that strains with many negative and positive chemical-genetic interactions are important for bioprocess-level predictions (**Supplementary Table 8**). Interestingly, and in accordance with recent findings regarding the differences in functional information encoded by negative vs. positive genetic interactions^14^, we found that negative chemical-genetic interactions were the primary driver of genetic interaction-based target predictions, and without them, the quality of the predictions was reduced substantially (See **Methods**, **Supplementary Table 9**).

In general, we found that compound bioactivity was correlated with our ability to make high-confidence predictions, as ~82% of compounds in our high confidence set inhibited growth >20% (**Supplementary Fig. 6**). However, the remaining ~18% of HCS compounds were associated with a more modest bioactivity (<20% growth inhibition), suggesting that even weakly bioactive compounds can yield functionally informative chemical-genetic profiles and that pre-screening for bioactivity may exclude some predictive profiles. A set of 296 compounds displayed extremely high bioactivity, with >90% growth inhibition, and nearly 60% (122) of these compounds were excluded from the final dataset because their interaction profiles did not meet thresholds for strain representation. Interestingly, chemical-genetic profiles for these highly bioactive compounds showed that mutants defective for two genes involved in amino acid transport, *GTR1* and *AVT5* were highly resistant and accounted for a majority of the read counts from these compound conditions. This suggests that these genes may play general roles in small molecule transport and that their deletions may confer general resistance to highly bioactive compounds (**Supplementary Fig. 7**), and we confirmed this finding for *gtr1*Δ cells in an independent experiment involving 23 different compounds (**Supplementary Table 10**).

### Defining the functional landscape of compound collections

To view the functional diversity of entire compound collections, each HCS compound was mapped onto the global genetic interaction profile similarity network at the location of the gene with the most similar genetic interaction profile to the compound’s top predicted biological process target^15,22^. The global network of genetic interaction profile similarities consists of 17 densely connected gene clusters, each representing a distinct biological process^14^ (**Fig. 3a**). The integration of the set of chemical-genetic profiles from a particular compound collection into the global genetic interaction profile similarity network allowed visualization of functional space covered by the compound collection (**Fig. 3b**) and enabled quantification of the diversity of targeted biological processes (**Fig. 4a**).

**Figure 3.**
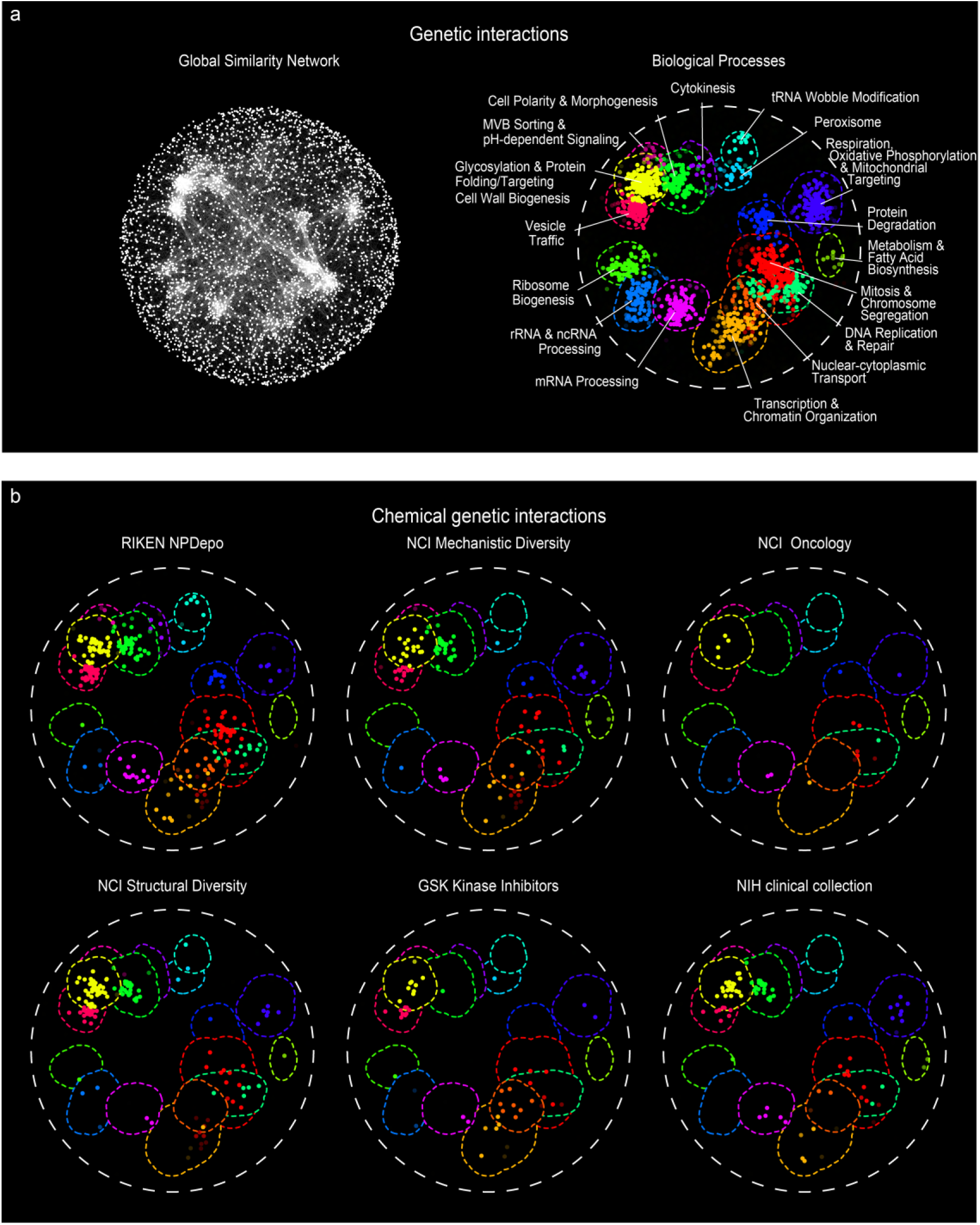
The functional landscape of diverse compound collections. (**a**). The global genetic interaction similarity network. (**a left panel**) Genes (nodes) that share similar genetic interaction profiles are connected by an edge in the global genetic interaction similarity network. Genes sharing highly similar patterns of genetic interactions are proximal to each other; less-similar genes are positioned further apart. (**a right panel**) Densely connected network clusters, color coded by functional enrichments annotations to 17 distinct biological processes. (**b**) Integrating genetic and chemical-genetic interaction profiles to predict biological processes targeted by HCS compounds. Colored nodes represent chemical compounds derived from the indicated collection. Each compound was placed on the map at the position of the gene with the most similar genetic interaction profile from the compound’s top predicted target process.

**Figure 4.**
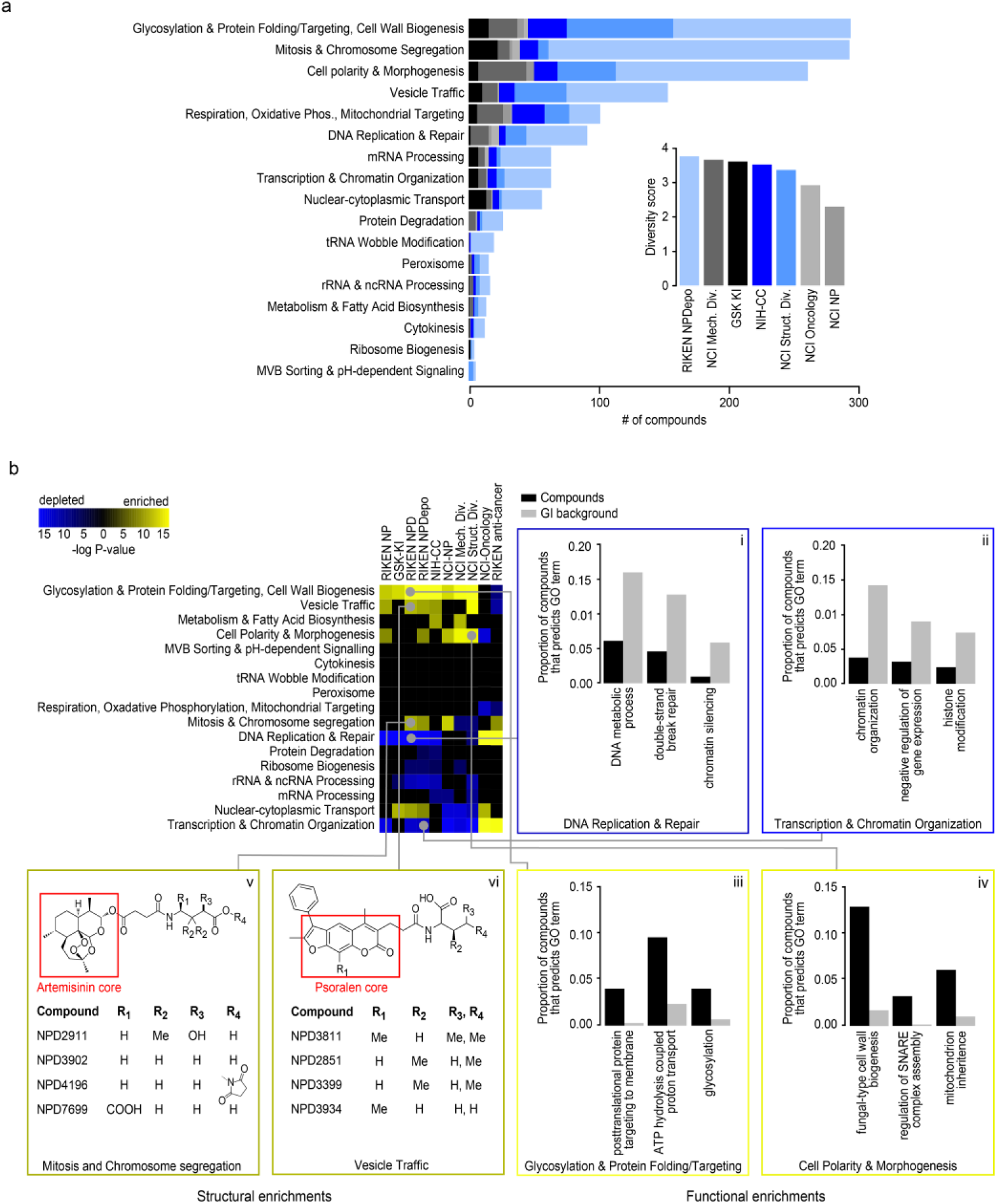
Functional signatures of compound collections. **(a)** Number of compounds within each collection’s HCS annotated to 17 distinct biological processes. **(inset)** Estimated functional diversity of each collection based on the uniqueness of chemical-genetic profiles from each library. **(b)** Compound collections and sub-collections were clustered based on their functional profiles. Collections whose chemical-genetic interaction profiles are enriched (yellow) or depleted (blue) for 17 distinct biological processes are shown. Sections are expanded **(i-vi)** to allow detailed visualization of significantly enriched GO biological process terms that drive the enrichment and depletion of target predictions, as well as enriched structural features of compounds predicted to target a biological process. Black bars represent the proportion of compounds within a collection annotated to a GO biological process, and grey bars represent the proportion of profiles in the GI background set annotated to the same GO term. (**v-vi**) Enriched structural features of artemisinin (**v**) and psoralen (**vi**) derivatives that are annotated to a specific biological process are presented with R-group decomposition.

Every major functional cluster in the genetic network appeared to be targeted by at least one compound screened in this study (**Fig. 3b**). However, glycosylation, mitosis, cell polarity, and vesicle traffic related functions were the most frequently targeted, suggesting that these bioprocesses are more susceptible to chemical perturbation in yeast (**Fig. 4a**). When corrected for the number of compounds, the RIKEN NPDepo collection was the most functionally diverse collection, whereas the NCI natural products collection (**Supplementary Fig. 8**) was the least diverse. The NPDepo library can be partitioned chemically and mechanistically into different subset collections, including natural products (NP), natural product derivatives (NPD), and anti-cancer compounds (a manually curated list of RIKEN compounds with known anticancer activity, **Supplementary Table 4**), all of which showed distinct functional signatures in terms of their targeted bioprocess predictions. Each compound collection targets a unique set of biological processes (**Fig. 3b and 4b**), suggesting that this global view of collection functionality can aid prioritization of screening efforts based on specific bioprocess targets of interest.

For the larger collections, we observed compounds targeting all 17 biological processes represented in the global genetic interaction similarity map. For example, the RIKEN NPDepo library was large and diverse enough to target all the major biological processes (**Fig. 4a-b**). Interestingly, the rate at which compounds targeted different biological processes differed from the distribution of genes across bioprocesses, suggesting a biased chemical target space (**Fig. 4b i-iv**). While each chemical library displayed a unique set of predicted bioprocess targets, common signatures emerged across several of the collections. For example, we observed a ~4-fold enrichment of compounds targeting glycosylation & protein folding related processes for most compound collections, including the NPDepo and NCI mechanistic diversity collections, which were designed to be relatively unbiased in terms of structure and functional annotations (**Fig. 4b**). Conversely, we saw a common depletion for compounds targeting DNA replication & repair and chromatin/transcription related processes, suggesting these processes are perturbed by compounds less frequently than expected, which could be an important consideration if, for example, targeting this biological process for cancer therapeutics is a major goal.

While enrichment for cytosolic targets and depletion for nuclear targets appeared as a general trend across several compound collections, exceptions were observed within specific libraries. In particular, for the NCI oncology collection, which is made up of anti-cancer agents largely directed towards the inhibition of cell division cycle functions and DNA replication/repair, we observed a strong enrichment for compounds targeting DNA replication & repair and transcription & chromatin organization relative to the expected background (**Fig. 4b,** p<0.001). The NCI oncology collection, along with the anti-cancer subset of the RIKEN collection, differed the most from the general trend observed for larger, less biased collections, reflecting the fact that these compounds have been selected for very specific purposes, which is largely confined to inhibiting growth of replicating cells.

The NIH-CC had a unique enrichment for compounds targeting metabolism and fatty acid biosynthesis, driven by GO predictions for sterol metabolic processes (**Fig 4b**, **Supplementary Table 11**). The majority of the compounds supporting this interact with cytochrome P450 enzymes^23–27^. In humans, compounds that inhibit or interact with cytochrome P450 have a high degree of drug-drug interactions^28,29^ In yeast, cytochrome P450 homologs ergosterol biosynthesis genes (*ERG11, ERG5, NCP1*). Thus, the yeast system provides a means of predicting compounds that interact with human cytochrome P450 enzymes, which could indicate compounds with a high degree of drug interactions.

The GSK kinase inhibitor (KI) library contains a characterized set of inhibitors of human kinases^30^. Three compounds from this collection were previously identified to bind human mitogen and stress activated kinases (MSK)^31^, and in yeast these had significant (p<0.05) enrichment for targeting the GO process of intracellular protein kinase cascade. This signaling pathway in yeast is mediated by the yeast mitogen activated protein kinase encoded by *SLT2,* the top single-gene target prediction for all 3 compounds (**Supplementary Table 12**), and has high homology to human ERK1, 2, and 4. Further, 5 compounds known to target human Polo-like kinase (PLK), were predicted in yeast to target the GO process nuclear import (**Supplementary Table 12**). The yeast homolog of PLK is *CDC5,* which is involved in regulating nuclear shape. These examples again suggest our yeast assay could be used to predict potential chemical bioprobes in human cells.

In general, the chemical-genetic functional signatures we observed appear to be related to cellular localization because cytoplasmic or cell surface related bioprocesses were more readily perturbed and thus enriched across diverse chemical libraries (p<0.0001), whereas nuclear processes were less susceptible to chemical perturbation, and compounds predicted to target these processes were depleted among many of the libraries tested (p<0.0001) (**Supplementary Table 13**). This may suggest that, in general, bioactive compounds are less likely to reach the nucleus, while cell surface and cytosolic targets may be more reactive. This is consistent with a previous study^32^, which reported that out of 1362 annotated drug targets with orthologs across 4 mammalian species, only 8.4% of these targets localized to the nucleus whereas 56% of targeted proteins localized to either membranes or the cytosol.

### Integrating structural and functional data

Because the RIKEN NPDepo contains sets of compound derivatives based upon variations of core scaffolds, we tested if compounds predicted to target similar functions were enriched for specific structural classes (**Fig. 4b v-vi**). Indeed, we found several instances where a large class of structural derivatives had similar predicted modes of action (**Supplementary Table 14**). For example, chemical-genetic profile similarity grouped a coherent set of artemisinin derivatives (**Fig. 4b v**) together within a broader subset of 358 compounds annotated to the “mitosis and chromosome segregation” biological process. While artemisinin is an effective anti-malarial drug, the cellular target(s) of this compound remain unclear^33^. In yeast, artemisinin is known to affect the cell cycle as well as mitochondrial function^34,35^. Furthermore, artemisinin has well-established effects on cancer cell cycle progression^36–38^. Our functional annotation supports both of these diverse roles for artemisinin because our artemisinin-related natural product (NP266) was annotated with 2 different biological process predictions: mitochondria cristae formation and microtubule cytoskeleton organization (**Supplementary Table 7**); however, the artemisinin derivatives that contain a relatively long side chain, extending from the three-ring core, have stronger predictions to a mitosis-related rather than a mitochondrial bioprocess-level target.

In another example, the furanocoumarin tricycle (psoralen) structural class is represented by multiple derivatives within the NPDepo library (**Fig. 4b vi**). Psoralen and its derivatives have been used to treat cutaneous T-cell carcinoma and dermatological conditions such as psoriasis and eczema^39^. The RIKEN NPDepo psoralen derivatives were frequently predicted to affect vesicle trafficking and membrane associated processes, and it is possible that other RIKEN NPDepo compounds with overlapping functional annotation could have a similar therapeutic potential.

### Targeted biological process validations and assessment of predictive power

In a previous study, the DNA content of yeast mutant strains harboring conditional alleles of essential genes were analyzed by flow cytometry, showing how each essential gene affects cell cycle progression and mapping specific cell cycle progression defects to different biological process^40^. For example, inhibiting the function of essential genes involved in translation causes an accumulation of cells in G1 phase (“G1” phenotype), reflecting insufficient protein synthetic capacity to transit the restriction point in G1 (referred to as Start in yeast), whereas inhibiting genes involved in DNA synthesis causes an accumulation of cells in S phase (“S” phenotype), and inhibiting mitosis genes results a G2 phase accumulation (“G2” phenotype). We performed high-throughput flow cytometry analysis on cell populations exposed to a set 67 different HCS compounds from the RIKEN NPDepo (**Supplementary Fig. 9**) that were predicted to cause specific cell cycle arrest phenotypes (**Fig. 5a-b**). In total, 27/67 (40%) of these compounds resulted in a cell cycle perturbation, and overall, 19/27 (70%) of compounds affecting cell cycle progression induced a phenotype consistent with our chemical-genetic predictions (**Supplementary Table 15**). For example, NPE94 was predicted to affect regulation of mitosis, and indeed, cells treated with this compound accumulated in G2 phase (**Fig. 5a**). Compounds displaying a cell cycle phenotype showed significant enrichment for each of the compounds’ predicted phenotypes over a background with permuted compound labels (G1: ~12-fold enrichment over background, p<0.001; G2: ~3-fold enrichment, p<0.01; S: 4-fold enrichment, p<0.001) (**Fig. 5a-b).** While only 40% of compounds induced a cell cycle phenotype in the single-dose and single time point tested, we suspect that an analysis using a dose curve and multiple time points would likely reveal cell cycle phenotypes among a number of the remaining ~60% of compounds predicted to affect this process.

**Figure 5.**
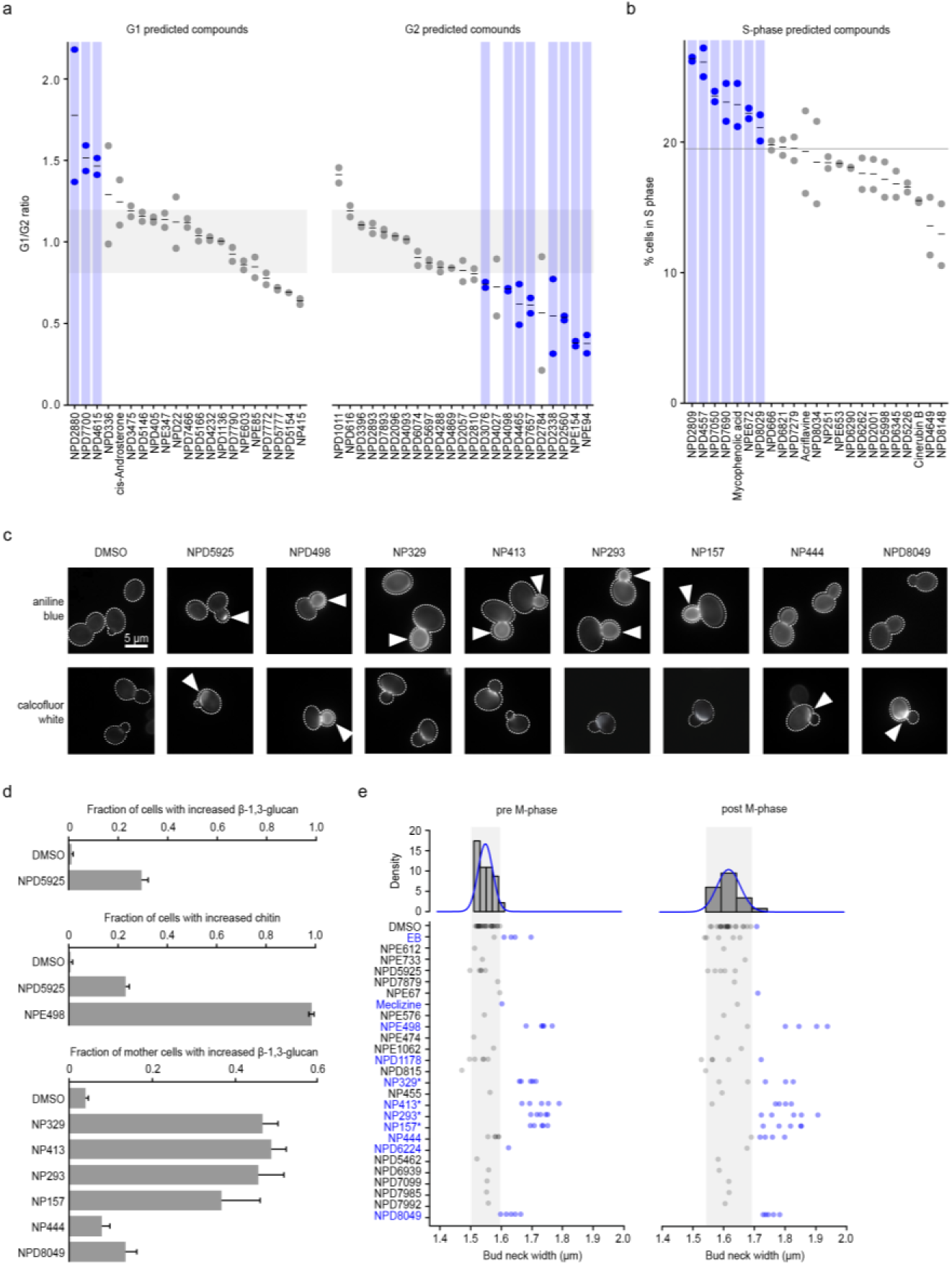
Large-scale validation of predicted target processes. **(a)** Comparison of observed and predicted cell cycle arrest phenotypes induced by 67 high-confidence compounds. Observed phenotypes were derived from flow cytometry analysis and predicted phenotypes were generated by mapping biological process annotations of the 67 compounds from this study to cell cycle arrest phenotypes via Yu et al. 2006^40^. Compounds that induced a G1 phase delay phenotype (G1/G2 ratio +1.5 standard deviations from the DMSO mean – above grey shaded box) or G2 phase delay phenotype (-1.5 standard deviations from the DMSO mean – below grey shaded box) are indicated (blue circles, n=2, biological replicates). (**b**) Compounds confirmed by flow cytometry analysis to cause defects in S phase progression (at least 1.5 standard deviations above the DMSO mean – above grey line) are indicated (blue circles, n=2 biological replicates). (**c**) β-1,3 glucan (AB=aniline blue) and chitin (CFW=calcofluor white) staining of cells treated with compounds predicted to affect the cell wall. Arrows indicate abnormal deposition of cell wall chitin or β-1,3 glucan. (**d**) Proportion of cells with increased β-1,3 glucan or chitin signal following treatment with predicted cell wall targeting compounds (n=3, mean ± S.E.). **(e)** Measurement of bud neck width in pre/post M-phase cells following treatment with 25 compounds predicted to target the cell wall (n=5). Blue text and circles indicate greater than average bud neck width. ^*^ denotes pseudojervine compounds.

As a second validation, we examined the activity of 25 compounds annotated to cell wall-related biological processes, utilizing several different cell biological readouts. To serve as controls, we selected 24 high-confidence compounds with equivalent growth inhibition and diverse bioprocess-level predictions but excluding “Cell Polarity and Morphogenesis” or “Glycosylation, Protein folding and Cell Wall Biosynthesis” bioprocesses predictions (**Supplementary Table 16**). Microscopic examination of fluorescent staining of two different cell wall polymers, β-1,3-glucan and chitin, revealed that 8/25 (32%) cell wall predicted compounds induced abnormal cell wall composition (**Fig. 5c-d**), and 10/25 (40%) caused increased bud neck width (**Fig. 5e**), a common phenotype of cell-wall-targeting agents^41,42^. Furthermore, 7 of these compounds caused hypersensitivity to zymolyase (**Supplementary Fig. 10a**), which degrades yeast cell wall β-1,3-glucan. In addition, 3/25 compounds caused rapid cell leakage similar to echinocandin B (**Supplementary Fig. 10b**), an antifungal drug that inhibits β-1,3-glucan biosynthesis. Among these compounds, we found a set compounds structurally similar to pseudojervines (**Supplementary Fig. 10c**). Based on this, we predicted, and confirmed that the poorly characterized parent compound jervine caused similar, abnormal glucan localization (**Supplementary Fig 10d**). The proportion of compounds that showed cell wall phenotypes in the cell wall-predicted set of compounds was significantly greater than that in the control compounds, even when all pseudojervines were treated as one compound. Overall, 48% (12/25) of the compounds predicted to target cell wall biosynthesis exhibited at least one cell wall defect associated phenotype, and 36% (9/25) of the compounds exhibited at least two phenotypes (**Supplementary Table 16**). In contrast, only 4% (1/24) of the control compounds showed any cell wall phenotypes (p < 0.05), (**Supplementary Table. 16**).

### Predicting compounds with dual targets

Our database of biological-process level annotation also offers the potential to screen for compounds that have multiple targets. Many pharmaceuticals perturb multiple cellular functions^43^, and identifying multifunctional compounds provides opportunities for drug repurposing and addressing potential side-effects of clinical agents^44^. We mined our HCS set of predictions to identify compounds that were associated with two, distinct biological processes (**Fig 6a**, **Supplementary Table 17)**. One of the top ranked compounds predicted to have multiple targets was NP214, a bleomycin A2 derivative. NP214 was predicted to target two different processes: (1) DNA replication (p<0.001) and (2) cellular proton transport (p<0.0001). The primary target of bleomycin and related compounds is DNA^45^; however, there is also evidence suggesting that these compounds perturb cellular membranes^46–48^, a secondary mode of action that could underlie bleomycin-induced side effect of lung fibrosis^46^. In mammalian cells, apart from its DNA activity, bleomycin has been shown to affect membrane redox potential and proton movement^49^. Moreover, bleomycin-iron complexes generate singlet oxygen and cause lipid peroxidation^50,51^. Thus, our chemical-genetic biological process predictions captured both the primary role of bleomycin (DNA damage) and secondary mechanisms that are consistent with known bleomycin side-effects.

**Figure 6.**
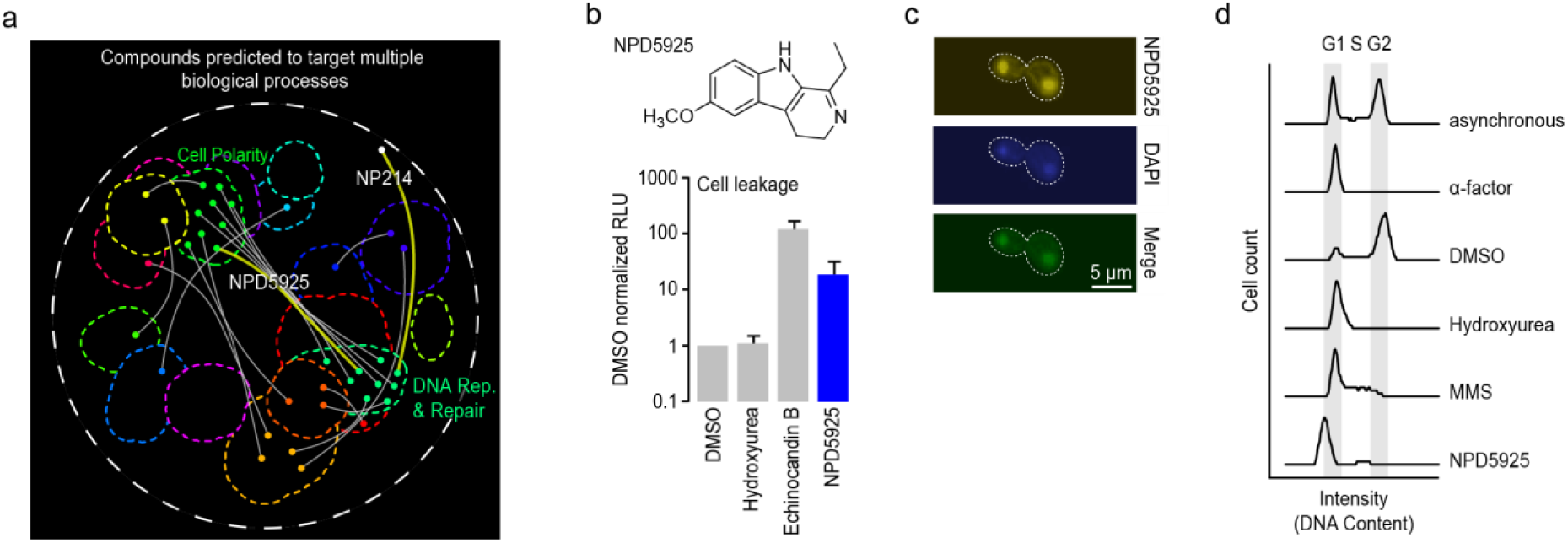
Identification of compounds with dual targets. **a)** Compounds predicted to target multiple distinct bioprocesses. Nodes indicate a predicted gene target located within a biological process-enriched network cluster defined in the global genetic interaction profile similarity network. Edges represent compounds predicted to target two distinct biological processes. NPD5925 was predicted to target the distinct processes of DNA catabolic process and fungal-type cell wall biogenesis (yellow edge). NP214 was predicted to target DNA replication and cellular proton transport (white node, yellow edge). **(b)** Measurement of cell leakage (adenylate kinase assay) from cells treated with DMSO, hydroxyurea, echinocandin B, and NPD5925 (n=3, mean ± S.E.). **(c)** Images of a cell stained with NPD5925 (fluorescent), DAPI, and the merged fluorescent signal. **(d)** Cell cycle analysis of cells following treatment with α-factor, DMSO, hydroxyurea (HU), MMS, and. NPD5925.

From a ranked list of dual target predictions (**Supplementary Table 17**) for HCS compounds, we observed a common coupling of DNA related processes and cell wall biogenesis (**Fig. 6a**). For example, when we exposed yeast cells to NPD5925, a novel RIKEN NPDepo compound that was predicted to affect both DNA catabolism (p<0.001) and cell wall biogenesis (p<0.001), they displayed cell surface defects, such as zymolyase sensitivity (**Supplementary Fig. 10**), and a cell leakage phenotype resembling that of echinocandin B (**Fig. 6b**). Because NPD5925 is fluorescent, we imaged its staining pattern and found that it localized to the nucleus, similar to DAPI (4',6-diamidino-2-phenylindole) (**Fig. 6c, Supplementary Fig. 11**); it also induced a G1/early S phase cell cycle arrest, similar to the arrest observed with high levels of hydroxyurea (**Fig. 6d**).^52^ While a compound that targets a pleiotropic gene could appear to perturb multiple, unique processes, we scanned the global yeast genetic interaction network for examples of genes displaying this type of genetic interaction profile; however, we were unable to find a single gene that could explain the dual bioprocess-level predictions of NPD5925 (**Supplementary Fig. 12**), which supports the dual functions of NPD5925, suggesting it perturbs both DNA catabolism and cell wall biogenesis processes independently.

Despite the clear dual target signal of these compounds, the effect of dose likely plays a significant role in separating the multiple modes of action of a compound. Indeed, a dose curve would likely help further dissect primary from secondary mechanisms of action of compounds. For example, in the case of NP214 and NPD5925, it is possible that DNA may be the primary target, and thus the DNA binding CG score signal would likely be apparent at lower doses, whereas the cellular proton transport or cell wall signals may only be detectable at higher doses. However, as we have screened dozens of DNA damaging agents that have not yielded these specific dual target signals, it is not likely that these findings are a consequence of general effects on DNA. While we still do not know the exact mechanism of NPD5925, we are able to deconstruct complex phenotypic consequences of a compound.

### New chemical genomic resources and analytical tools

We generated an active database named MOSIAC (http://mosaic.cs.umn.edu/), housing all our chemical-genetic screens. The MOSAIC database catalogs the structural and basic physical properties of all compounds tested, including their bioactivity, chemical-genetic profiles, as well as the biological process and gene-level target predictions. We also developed novel software tools, called BEAN-counter (Barcoded Experiment Analysis from Next-generation sequencing), for processing raw sequencing data into chemical-genetic interaction profiles, and CG-TARGET (Chemical-Genetic Translation via A Reference Genetic interaction nETwork) for predicting biological process-level targets from chemical-genetic interaction profiles. These new software tools are available at http://github.com/csbio/.

### The Bioprocess Diversity Set: A collection of functionally characterized bioactive compounds

We distilled the most functionally diverse compounds from all 7 libraries analyzed in this study into a new “Bioprocess Diversity Set” (**Supplementary Table 18**), which represents a selected collection of our HCS bioactive compounds whose targets span the functional landscape of the cell. We also selected “Bioprocess Specific Sets”, each consisting of a set of compounds predicted to target one of the 17 different biological processes represented in the global genetic interaction profile similarity network (**Supplementary Table 19**). We anticipate that these new compound collections should provide a powerful new resource for modulating cellular physiology through diverse perturbations and streamlining the chemical-genetic discovery pipeline, enabling a focused analysis on specific biological processes of interest. The Bioprocess Diversity Set and the Bioprocess Specific Sets can easily be sorted to focus on individual compound libraries, including the NCI, NIH, GSK and RIKEN NPDepo libraries.

## DISCUSSION

Our high-throughput chemical-genetics platform addresses a need for an unbiased, whole cell method that provides rapid, functional annotation of compound libraries. We used this system to screen 13,524 compounds across 7 different libraries, yielding rich chemical-genetic profiles and high-confidence functional predictions for a set of 1522 compounds. We cataloged the complete dataset as an open chemical-genetics resource (http://mosaic.cs.umn.edu/).

Our functional annotation of chemical libraries offers a strategy for prioritizing compounds that display bioactivity directed towards particular biological processes. The scale of functional annotation also provides a global view of the chemical activity within a library, which should allow testing of general hypotheses relevant to chemical biology. Importantly, the high-throughput nature of this assay provides opportunities for systematic, large-scale functional analysis of natural extract collections. Natural extract collections are often far more expansive than pure compound libraries and may contain broader mechanistic diversity. Functional annotation of these collections would help identify and prioritize promising extracts for detailed fractionation^10^.

Our approach highlights the use of drug hypersensitive and diagnostic mutant sets for compound characterization, which allowed us to interrogate more compounds and use smaller quantities. Certain drug efflux transporters can be dedicated to certain classes of drugs, such as *PDR5* which has a documented specificity to steroid drugs^53^. Thus, although we only explored one genetic background for sensitization, it is possible to construct new yeast mutant collections using different genetic backgrounds tailored specifically for hypersensitivity to particular drug classes. In addition, while we selected a diagnostic pool of mutants specifically for genome-wide functional annotation, diagnostic pools with specific functional biases could be designed to investigate particular cellular processes or targets. Moreover, the diagnostic pool may be further reduced in size for greater multiplexing, as we found as few as 157 strains had equivalent predictive power as the entire non-essential collection of ~4900 strains (see **Methods**).

One advantage of our approach is that we can functionally characterize compounds that do not show strong bioactivity. While bioactivity was predictive of our ability to make high-confidence predictions, it was not absolutely necessary. Pre-screening for bioactivity, which is a common approach^11,21^ can potentially exclude compounds with specific but possibly nonessential modes of action. For example, ~18% (270 of 1518) of the HCS compounds we identified for which we had bioactivity measures inhibited growth <20%. Indeed, weakly acting compounds targeting specific functions represent a starting point for chemical modifications to improve bioactivity.

Biological process target predictions derived from the global yeast genetic interaction network provides a roadmap, not only for other microorganisms (e.g. *S. pombe, E. coli*), but also for mammalian systems. Importantly, the construction of genetic interaction maps in human cell lines is possible, as is the mapping of chemical-genetic interactions^54–56^. Thus, the same approaches and predictive tools we implemented in yeast can be adapted and applied as a general strategy to map analogous chemical-genetic networks for human cells. More generally, combinatorial genetic and chemical-genetic approaches can be used to identify new drug leads that work synergistically to expand our understanding of druggable target space^3,57^.

## Acknowledgements

This work was supported by RIKEN Strategic Programs for R&D. J.S.P. and S.C.L were funded by a RIKEN Foreign Postdoctoral Fellowship. S.W.S. is supported by an NSF Graduate Research Fellowship (00039202), an NIH Biotechnology training grant (T32GM008347), and a one-year BICB fellowship from the University of Minnesota. H.O. is a research fellow of the Japan Society for the Promotion of Science. R.D., J.N., E.W., and C.L.M. are supported by National Institutes of Health Grants 1R01HG005084-01A1, 1R01GM104975-01, and R01HG005853 and National Science Foundation Grant DBI 0953881. C.B. and Y.O. are supported by JSPS KAKENHI Grant Numbers 15H04483. C.B. and B.A. were supported by the Canadian Institutes of Health Research, FDN-143264 and FDN-143265, respectively. C.L.M, C.B., M.C., J.L., and B.A. are supported by the Canadian Institute for Advanced Research Genetic Networks Program. Y.O. is supported by Ministry of Education, Culture, Sports, Science and Technology, Japan Grant for Scientific Research 24370002 and JSPS KAKENHI Grant Numbers 15H04402. A.B. is supported by a Lewis Sigler fellowship at Princeton University. G.W.B. and N.P.T are supported by Canadian Cancer Society Research Institute impact grant 702310. K.S. is supported by CREST from JST. We thank Astellas Pharma Inc. (Tokyo, Japan) for their kind gift of Micafungin. We thank Tamio Saito for help with NPDepo compound access. Sequencing was provided by RIKEN Center for Life Science Technologies, Division of Genomic Technologies, Genome Network Analysis Support Facility (GeNAS) RIKEN CLST and the University of Chicago.

## Author contributions

C.B., M.Y., J.P. and S.L. conceived the project. J.P., S.L., R.D., and S.S. designed the chemical genomic screens. J.P., S.L., J.B., R.O. M.Y., Y.Y., E. D-A. performed the chemical genomic experiments. C.M., R.D., S.S., J.N., H.S., and E.W. designed the analysis software and performed analysis. G.B., N.T., and S. L. performed cell cycle experiments. J.P., K.A., M.L., and A. B. designed the sensitized yeast. Y.O., H.O., A.G., K.K. performed validation of cell wall targeting compounds. R.D. and M.C. designed the diagnostic mutant collection. H.O. and H.H. provided and curated NPDepo compounds. K.S. provided sequencing and analysis. J.V. edited the figures of the manuscript. The manuscript was written by J.P., S.L., C.M., and C.B. with input and editing from all authors.

## Online Methods

### Constructing a genome-wide drug sensitive yeast deletion collection

Construction of the *pdr1∆ pdr3∆ snq2∆* triple mutant is described in Andrusiak 2012^19^. Briefly, *PDR1* was deleted in the SGA query strain (Y7092)^58^ by replacement with the *natMX* antibiotic resistance marker, which provides resistance to the drug nourseothricin (NAT). To construct the *pdr1∆ pdr3∆* double mutant, *PDR3* was deleted in the *pdr1∆* mutant by replacement with the *K. lactis URA3* autotrophic marker, which permits cells to grow on synthetic media lacking uracil. The *pdr1*∆, *pdr3*∆, and *snq2∆* single or double mutants were constructed by replacing the wild type gene with the *natMX, K. lactis URA3,* and *K. lactis LEU2* markers, respectively. The *natMX, Kl.URA3* and *Kl.LEU2* markers were amplified from plasmids using primers designed with 50 base pairs of sequence homologous to regions upstream and downstream of the genes. PCR amplicons were transformed into the appropriate strains using lithium acetate and polyethylene glycol-based transformations^59^. Deletion of the native gene and integration of the marker at the correct locus was confirmed using a series of PCR-based confirmations. Confirmation primers were designed specific to regions both flanking the integration site and internal to the inserted marker to interrogate both the full length of the inserted marker and the 5’ and 3’ boundaries.

The MATα *pdr1∆::natMX pdr3∆::KI.URA3 snq2∆::KI.LEU2* (y13206) query strain carried the *can1∆::STEpr-SP_his5* and *lyp∆* SGA reporters. *STEpr-SP_his5* is an auxotrophic marker that allows only *MATa* cells to grow in the absence of histidine, while the *can1∆* and *lyp∆* deletions allow haploid cells to grow in the presence of the drugs canavanine and thialysine, respectively. The *MAT*α query strain was crossed to an ordered array of *MAT****a*** *xxx∆::kanMX* deletion mutants and the resulting heterozygous diploids were transferred to media with reduced carbon and nitrogen to induce sporulation and the formation of haploid meiotic progeny. The resulting spores were transferred to synthetic media lacking histidine and containing canavanine and thialysine to select for the *MAT****a*** meiotic progeny. Cells were then transferred to synthetic media lacking uracil and containing NAT to select for growth of cells carrying both the *pdr3∆*::KI.*URA3* and *pdr1∆::natMX* deletions. Finally, these cells were transferred to synthetic media lacking uracil & leucine and containing G418 & NAT to select for the desired *pdr1∆ pdr3∆ snq2∆ xxx∆* mutants. This protocol was adapted from Kuzmin et al 2016^60^.

### Assessing compound hit rate of sensitized yeast strains

The chemical sensitivity of deletion mutants was assessed using a high-throughput chemical growth inhibition halo assay. After growing WT, *pdr1∆ pdr3∆* and *pdr1∆ pdr3∆ snq1∆* mutant yeast strains overnight to saturation, cultures were standardized to an OD600 = 4.0 and 2 mL was added to a 50 mL stock of 2% YP (10 g/L yeast extract, 20 g/L peptone) + 2% galactose + 1% agar (YPGal). Seeded plates were prepared by pouring 10 mL of culture into NUNC square plates and drying for 10 minutes to facilitate compound absorption. Robotic pinning with the Biotec ADS384 was used to transfer 0.2 μL of each natural product to the seeded plates at a density of 88 compounds per plate; 440 diverse compounds (**Supplementary Table 1**) from the RIKEN NPDepo were evaluated in total. After incubating for 24 hours at 30 °C, plates were imaged and the visible areas of growth inhibition were measured using JMicrovision (Version 1.2.2. http://www.jmicrovision.com). A compound was deemed toxic if it generated an area of growth inhibition with a diameter greater than 1 mm. Thus, we assessed the number of compounds that perturbed growth (e.g. compound hits) of WT, *pdr1∆ pdr3∆* and *pdr1∆ pdr3∆ snq1∆* mutant strains.

The chemical-sensitivities of the top drug-sensitive deletion mutants identified from the adapted assay were confirmed by growing deletion strains in the presence of the tested drug (34.4 μΜ benomyl, 25 nM micafungin, or 1% DMSO) for 24 hours and recording the resulting optical density at 600 nm. Strains tested harbored deletions either in a wild-type background or in the drug-hypersensitive *pdr1∆ pdr3∆ snq2∆* background. Values plotted are percentages calculated by dividing the OD600 measured after growth in DMSO by the OD600 measured after growth in the specific concentration of compound and multiplying by 100 (**Fig 1. c-d**). Y7092^58^ was used as the WT control and the *pdr1∆ pdr3∆ snq2∆* mutant was used as the drug hypersensitive control. (n = 3).

### Defining the diagnostic gene set for optimized chemical genomic screens

A diagnostic set of 310 genes was selected by combining the output from two methods: a computational strategy and a manual selection. A set of 157 genes was selected by identifying functionally relevant genes using a computational approach called COMPRESS-GI (Deshpande et al. *in preparation*). Because genetic interaction profile similarity can be accurately measured using only a subset of the genome-wide profile, the COMPRESS-GI method selects genes to be included in a genetic interaction (and chemical-genetic) profile to maximize the agreement between pairwise gene similarities computed from the compressed profile and gene co-annotation information from the Gene Ontology. Selection of such a subset of genes is useful for our chemical genomics study because the reduced chemical-genetic profile for each compound is directly compared with the corresponding reduced genetic interaction profiles, which generates accurate compound-gene similarities based on a small set of mutants. The COMPRESS-GI algorithm is described and evaluated in depth elsewhere (Deshpande et al. *in preparation*).

In addition to the 157 genes selected with the computational approach, we also manually selected 236 genes. The logic for the manual method was to pick any single member of the same pathway/complex because members of the same pathway/complex possess similar genetic interactions. Hence, picking one gene from each pathway/complex should be sufficient to cover the genetic network space associated with all the genes in that pathway/complex. We applied 2-dimensional hierarchical clustering to cluster gene deletion mutants based on their genetic interaction profiles, and then manually selected strains that displayed rich genetic interaction profiles representative of each of the 17 functionally enriched cluster from the global genetic interaction profile similarity network (Costanzo et al., 2016^14^) to generate a minimal subset of yeast deletion mutants that re-capitulated the majority of functional profiles observed in our reference map.

Both the COMPRESS-GI and manual gene selection methods were applied using a filtered, non-essential yeast genetic interaction dataset^15^ where strains observed to exhibit extreme read counts in barcode sequence (top/bottom 10%) were removed. Also, in cases where multiple different mutant alleles were available for the same gene, the allele with the highest number of genetic interactions (highest interaction degree) in its genetic interaction profile was chosen. We found 83 genes in common between the computational and manually-derived lists, suggesting that the two methods had good agreement with respect to which genes were informative. The union of genes from the two selection methods comprised the initial diagnostic strain set (**Supplementary Table 1**).

Pilot experiments using this diagnostic set (**Supplementary Table 1**, diagnostic pool version 1) revealed a number of mutants that still exhibited abnormally high or low barcode counts in all experiments. These were removed to generate a collection of 310 strains for the final version of the diagnostic strain set (**Supplementary Table 1**, diagnostic pool version 2).

### Optimization of signal detection/sequencing parameters

Initial optimizations were conducted using a preliminary diagnostic pool of 491 strains. This pool of deletion mutants was constructed by pinning frozen 96-well glycerol stocks of each strain onto Nunc Omni Tray plates containing YPD + G418 solid media and incubating for 2 days at 30°C. Each plate was then flooded with 10 mL of YPD liquid media and a cell spreader was used to re-suspend grown colonies. The resulting cell suspensions were transferred to a 50 mL conical tube where glycerol was added to a 15% final concentration. Finally, the pool was adjusted to a final concentration of 50 OD600/mL by dilution or centrifugation and stored at −80 °C until required. To assay the mutant pool for drug-hypersensitivity, cells were thawed, counted using a haemocytometer, and diluted to seven different final inoculum densities (3727-58 cells / strain) in YP + 2% galactose in a 96-well flat-bottom plate. Cultures were then spiked with either 34.4 μΜ benomyl, 25 nM micafungin, or a 1% DMSO control. After growing for 18, 24, or 48 h at 30 °C, cells from each well were harvested by centrifugation. Genomic DNA was purified from the harvested cells by re-suspending in 125 μL of zymolyase buffer (1 mg/mL) and using the QIAextractor (Qiagen) as per manufacturer's instructions, with a 100 μL elution volume.

Barcodes were amplified from each of the wells using multiplex primers as described elsewhere^61^ for 20, 25, or 30 cycles. Samples were gel purified from 2% agarose and assessed for quality using the Kapa Illumina qPCR kit. Samples were sequenced at a loading concentration of 10 pM on an Illumina HiSeq2000 as a single uninterrupted read (“read through”). The 30 cycle samples were also sequenced using a “separated read” strategy, where the barcodes were read in a first sequencing step, while the multiplex tags were read after a second priming step. Output from “read-through” and “separated read” runs were then compared. The signal to noise was calculated by taking the mean CG score of the top 10 array genes divided by the standard deviation of all array genes CG scores. This was done for each drug, PCR cycle, cell density, culture combination.

### Multiplex tag design and 768-plex primer selection

We designed one thousand 10 bp multiplex tags such that (1) the Levenshtein distance between any two tags was greater than 3, and (2) the tags were balanced in terms of nucleotide distribution. Condition (1) ensures that multiplex tags are maximally distinguishable even with a small number of sequencing errors while condition (2) ensures that the GC content and predicted melting point of all tags were within a small range. Because the space of multiplex tags is too large to exhaustively enumerate, we generated random multiplex tags and selected tags iteratively if both conditions were true. Primers containing the Illumina sequencing adapter, common priming site for the UPTAG barcode, and 1000 selected 10 bp multiplex tags were synthesized (Sigma, St. Louis, MO, USA), arrayed in 96-well plates. To assess amplification performance of the multiplex tags, we performed 1000 identical pooled growth experiments on the diagnostic strain pool under control conditions (DMSO). Samples were processed as described above and sequenced on an Illumina MiSeq lane (1000-plex). We used the count distribution to identify 8 plates (768 multiplex tags) with the most uniform distribution of read counts (**Supplementary Fig. 13**), and discarded plates containing multiplex tags with highly divergent reads counts. These 8 plates of multiplex tags with equivalent performance were used in all subsequent experiments (**Supplementary Table 3**).

To test the effects of multiplexing on the chemical genetic interaction signal, we selected a set of 768 compound conditions, including DMSO controls, known agents, and novel bioactive compounds from the RIKEN collection. For each assay we used the optimal pooled growth conditions defined above. We included a subset of compounds also screened in the Parsons *et al.* 2006 dataset as controls at every plexing level (96, 192, 384, and 768)^10^. We dosed the pooled cells at a level that inhibited growth by 20-50% compared to the DMSO control. Genomic DNA extraction, PCR, sample prep, and sequencing were performed as described above.

### Screening the NPDepo/NCI/NIH/GSK collections

We performed our pooled growth assay with the diagnostic mutant collection under optimized conditions as described above. Excluding controls compounds, we performed two screens totaling 13524 conditions, which represented 13431 uniquely-named compounds. In the initial batch of compounds examined, we screened the first 9840 members of the growing RIKEN NPDepo, and in the second batch, we screened six publicly available plated libraries: the NCI Natural Product (117 compounds), Approved Oncology (101), Structural Diversity (1599), and Mechanistic Diversity (821) collections, the NIH Clinical Collection (720), and the GlaxoSmithKline kinase inhibitor collection (326). The NPDepo is maintained as 1 mg/mL stocks, and we screened it at a final concentration of 10 μg/mL, with the exception of a number of compounds that received additional lower dosing in a pilot experiment (**Supplementary Table 4**). All remaining collections were screened at 100 μΜ, with the exception of the NCI Mechanistic Diversity set (10 μΜ) (**Supplementary Table 4**). Selected compounds were re-screened at lower concentrations if the initial concentration resulted in severe growth inhibition. The diagnostic mutant pool was grown in 200 μL cultures in 96-well plates. Each plate had 88 test compounds, 4 control compounds, and 4 internal DMSO conditions, (**Supplementary Fig. 14)**. Each lane consisted of 7 compound plates and one DMSO control plate, and every plate had 3 independent PCR replicates. For pairs of replicates of our control compounds, we measured Pearson correlation coefficients of 0.94, 0.95, 0.93, and 0.92 for our control compounds, respectively (Benomyl, Micafungin, MMS, Bortezomib). Thus, 3 replicates were sufficient to ensure high-quality, quantitative chemical genomic profiles. The primer set used to amplify each plate was shuffled for each replicate in such a way that each compound replicate would not use any single multiplex tag more than once. The primer set used to amplify the DMSO plate was different for each lane. The control compounds give very distinct CG profiles and were used to ensure proper plate orientation at all steps of the process. Culture OD was measured at 0, 24, and 48 h, and growth at 24 h relative to the DMSO control was used as a measure for bioactivity.

Following growth, genomic DNA was extracted as described above. The genomic extractions for each plate were amplified in triplicate using three unique multiplex primer plates (3 technical replicates). We used 768-plexing per lane, which means each sequencing lane contained PCR amplified barcodes from eight 96-well plates. We ensured each of the multiplex primer plates were used to amplify the DMSO plates allowing us to detect and remedy any potential multiplex primer biases following sequencing. Following PCR, samples were pooled first by plate, then by lane. The “per lane” samples were purified by *2%* agarose gel and the product quantified by qPCR as described above. All samples were run at a loading concentration of 10 pM as single-end, 50 bp reads on an Illumina Hiseq2000.

### Description of compound collections

#### RIKEN NPDepo

The RIKEN Natural Products Depository (NPDepo) is a public depository of small molecules. Currently, the NPDepo chemical library contains 39,200 pure compounds, half of which are natural products and their derivatives^62^.

Each of the remaining collections are publicly available and can be requested at the sites listed below.

#### NIH-Clinical collection

http://nihsmr.evotec.com/evotec/sets/ncc

#### NCI-Structural diversity collection

https://www.dtp.nci.nih.gov/branches/dscb/div2_explanation.html

#### NCI-Mechanistic diversity collection

https://www.dtp.nci.nih.gov/branches/dscb/mechanisticII_explanation.html

#### NCI-Oncology collection

https://www.dtp.nci.nih.gov/branches/dscb/oncology_drugset_explanation.html

#### NCI-Natural products collection

https://www.dtp.nci.nih.gov/branches/dscb/natprod_explanation2.html

#### GSK-Kinase inhibitor collection

https://www.ebi.ac.uk/chembldb/extra/PKIS/

### Computing molecular descriptors for all screened compounds

SMILES and InChI string representations of all molecules were generated using the OpenBabel cheminformatics toolkit^63^ (http://openbabel.org) and its python wrapper, pybel^64^. All molecular descriptors (column J through the last column) were calculated using PaDEL-Descriptor^65^, a wrapper for the Chemistry Development Kit cheminformatics toolkit^66^.

### Chemical-genetic interaction scoring method

The first step in the chemical genetic interaction scoring pipeline was to count the number of reads that mapped to each combination of knockout strain and chemical condition. All 50 bp sequencing reads were collected from the sequencing instrument as fastq files. All sequences in the fastq files containing the common primer sequence (U1 primer) were retained. No more than two errors were allowed (command: agrep −2) when matching the common primer sequence. Each retained sequence was then split into a “multiplex tag” (bases 1-10) and a “barcode” (bases 28-47). Each multiplex tag was matched against the list of expected multiplex tags, with no errors allowed for a match. Each barcode was matched to the barcodes in the diagnostic strains, with two errors allowed. A double hash data structure (one hash for the multiplex tag and a second for the barcode) was used to record the number of reads that matched to a particular combination of multiplex tag (identifies the condition) and barcode (identifies the strain).

Multiple filters were applied to clean the read count data before further processing. First, data from the *gtr1*Δ and *aνt5*Δ strains were removed, because these strains were over represented following growth in a specific set of conditions and thus accounted for a large majority of the read counts (**Supplementary Fig. 7**). Conditions that possessed low read counts across specified number of strains (see parameter settings below) were then removed (“condition count filter”), followed by strains with low read counts across a certain number of conditions (“strain count filter”). Details on each filter and its application are found in the “Applying the chemical-genomics” section below. The read count total for each strain-condition combination was then transformed into log space or set to NaN if 0.

Chemical-genetic interactions were then calculated by comparing the profile of log read counts from each condition across all strains to a reference profile of log read counts across all knockout strains. First, the reference profile was calculated as the log of the strain-wise average profile across all DMSO control read count profiles (excluding read counts less than 20, which were set to NaN and excluded from the calculation). Second, the log read count profile for each condition was LOWESS normalized with respect to the reference DMSO profile (span = 30%). Finally, chemical-genetic interactions were computed as z-scores representing the standardized deviation of each strain in the condition profile with respect to its counterpart strain in the reference profile.

Chemical-genetic z-scores were calculated in two different ways: the first using a condition-specific standard deviation and the second using a reference standard deviation vector. The condition-specific standard deviation was calculated on the middle 75% of the strain-wise deviations between each condition profile and the reference profile. The reference standard deviation vector was calculated using all DMSO control profiles, as the square root of the LOWESS-derived estimate (span = 30%) of the squared deviations with respect to the reference profile. One reference standard deviation vector was computed for the positive deviations and another for the negative deviations; the final reference standard deviation vector was specific to each individual condition depending on the signs of the deviations in that condition. Chemical genetic z-scores were then calculated as the deviation from the reference divided by the larger of 1) the condition-specific standard deviation or 2) the appropriate value from the reference standard deviation vector (pos. or neg. standard deviation). Positive z-scores suggest resistance to the condition, and negative z-scores suggest sensitivity to the condition relative to the fitness of each mutant in the control condition.

We observed a multiplex tag batch effect in the z-score dataset, wherein the average correlation between conditions with the same multiplex tag was larger than the average correlation between conditions with different multiplex tags. To address this effect, we first removed the multiplex tags with the worst batch effects (“batch effect filter”). Then, we applied Fisher’s linear discriminant analysis (LDA) to reduce the pairwise condition correlations observed within multiplex tags vs. between multiplex tags, a method which has been previously used to remove batch effects from genetic interaction screens^58^. LDA components were removed based on PR curve analysis of the multiplex tag effect (“multiplex tag effect parameters”, see settings for each dataset below). To prepare the data for LDA, duplicated conditions with the same multiplex tag were removed such that only one compound replicate was retained per multiplex tag.

Following the removal of the multiplex tag effect, we performed two final processing steps to yield the final dataset. In the “SVD” step, a large technical artifact was removed using singular value decomposition (SVD). This effect was deemed to be technical, as it was a general signature that occurred in ~1/3 of the condition profiles and had only a small number of weak Gene Ontology biological process enrichments. When removed (by removing projections of compounds’ profiles onto the first SVD-derived component), correlations with genetic interaction profiles were not noticeably affected. In the “replicate collapsing” step, profiles from technical replicate conditions were collapsed into single, mean profiles. To determine which replicates would be collapsed into each final profile, a graph was constructed in which nodes were the technical replicates and edges indicated a correlation between two replicates above a certain threshold (“replicate correlation threshold”).

### Chemical genetic interaction scoring algorithm

~~~
Pseudocode for the scoring algorithm is included below.
*x_i_* = read count profile for condition *i* across all strains
*x_log_i_* = log(*x_i_*)
*i_reference* = indices of reference conditions (DMSO control profiles)
*x_ref* = log(strain-wise mean vector across all *x_i_* for *i* in *i_reference,* excluding values in each *x_i_* of less than 20 read counts)
#Compute continuous estimates of positive and negative standard deviations across the reference profile (compute lowess on squared deviations)
*X_dev_pos, x_ref_pos, x_dev_neg, x_ref_neg* = empty vector
For each condition index *i_ref* in *i_reference*:
     *x_norm_i_ref_* = *x_log_i_ref_,* lowess-normalized with respect to *x_ref*
     *x_dev_i_ref_* = *x_norm_i_ref_*^_^*x_ref*
     *x_dev_sq_i_ref_* = (*x dev_i_ref_*) ^^^ 2
     For *j* in 1:length(*x*_*ref*):
          if (*x_dev*)*j* >= 0:
               Append (*x_dev_sq_i_ref_*)*j* to *x_dev_pos*
               Append *x_refj* to *x_ref_pos*
          if(*x_devi*)*j* < 0:
               Append (*x _dev_sq_i_ref_*)to *x_dev_neg*
               Append *x_refj* to *x_ref_neg*
     #Calculate continuous standard deviations
     *x_std_pos* = *x_dev_pos,* lowess-normalized with respect to *x_ref_pos*; one value is retained per element in *x_ref,* yielding a vector of standard deviations that each correspond to one strain in the *x_ref* profile
     *x_std_neg* = same as *x_std_pos,* but derived from *x_dev_neg*
# Compute chemical-genetic interaction scores for all condition profiles
For each condition index *i*:
     *x_norm_i_* = *x_logi,* lowess-normalized with respect to *x_ref_*
     # Deviation from the reference profile
     *x_dev_i_* = *x_normi* – *x_ref*
     # Condition-specific standard deviation for condition *x_i_*
     *std_constant_i_* = standard deviation across middle 75% of *x_dev_i_*
     #Reference standard deviation for condition *x_i_*
     *Std_continuous_i_* = empty vector
     For *j* in 1:length(*x*_*devi*):
          if (*x_dev_i_*)*j* >= 0:
               (*std_continuous_i_*)*j* = *x_std_pos_j_*
          else:
               (*std_continuous_i_*)*j* = *x_std_neg_j_*
     # The maximum standard deviation applicable to each element in *x_dev_i_*
     *Std_max_i_* = max((*std_constant_i_*)*_j_*, (*std_continuousi*)_*j*_) for each element with index *j* in *x_dev_i_*
     # Chemical-genetic z-score profile
     *z_i_* = *x_dev_i_*/*std_maxi*
~~~

### Applying the chemical genetic interaction scoring method to chemical genomic screens in this study

Data in this study were collected from two large screening efforts. The “RIKEN” screen encompasses all compounds originating from the RIKEN Natural Product Depository, and the “NCI/NIH/GSK” screen encompasses all remaining compounds (the NCI collections, NIH Clinical Collection, and GSK Kinase Inhibitor collection). The same basic experimental and computational procedures described above were used with a few minor variations, which are documented below.

Filters:

- RIKEN screen

- Condition count filter: The condition was removed if it had < 200 total read counts across all strains.
- Strain count filter: No filter was applied
- Batch effect filter: A multiplex tag, and its corresponding conditions, was removed if the average pairwise correlation of its conditions was 0.4 or higher. The average correlation value was calculated using only DMSO control profiles.
- NCI/NIH/GSK screen

- Condition count filter: For each condition to pass this filter, 50% of strains in the condition vector were required to possess more than 20 read counts.
- Strain count filter: For each strain to pass this filter, 50% of the conditions in the strain vector were required to possess more than 20 read counts. Batch effect filter: A multiplex tag, and its associated conditions, was removed if the average pairwise correlation of its conditions was 0.4 or higher. The average correlation value was calculated using all profiles from the screen.

Multiplex tag effect parameters

- RIKEN screen: 6 LDA components were removed
- NCI/NIH/GSK screen: 5 LDA components were removed

Order of SVD and replicate collapsing steps

- RIKEN screen:

1. SVD removal of large technical effect
2. Replicate collapsing (replicate correlation threshold: 0.7)
- NCI/NIH/GSK screen:

1. Replicate collapsing (replicate correlation threshold: 0.5)
2. SVD removal of large technical effect

All chemical-genetic interaction heat maps were hierarchically-clustered on both axes using 1 – *S_C_*(*X, Y*) as the distance measure with average linkage (where *X* and *Y* are chemical-genetic interaction profiles and *S_C_* is the cosine similarity between two vectors,

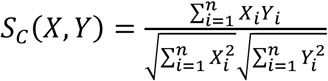
They were visualized using Java TreeView.

### Predicting compounds’ modes of action based on chemical-genetic and genetic interaction profiles

#### Genetic interaction dataset

Genetic interaction data was obtained from a recently assembled *S. cerevisiae* genetic interaction dataset^14^. The genetic interactions were derived from most recent quantitative fitness observations of non-essential, double mutant strains described in Costanzo et al., 2010, and Costanzo et al., 2016)^14,15^. The data consist of a genetic interaction score (the difference between each double mutant’s observed and its expected fitness values) and an associated p-value for each double mutant. The data were preprocessed by setting all epsilon scores to zero for which the associated p-value was greater than 0.05.

From this preprocessed dataset, we identified a subset of 1505 high-signal genetic targets that would provide a robust basis for target prediction from our chemical-genetic data. The criteria for selecting each query GI profile were: 1) it must have greater than 40 significant interactions (degree > 40), 2) the sum of its cosine similarity scores with all other query profiles must be greater than 2, and 3) the sum of its dot products with all other query profiles must be greater than 2. These criteria helped to define a set of genetic interaction screens with sufficient signal for correlation with chemical genetic profiles. The final genetic interaction dataset consisted of the 1505 query GI profiles that passed these criteria, each of which reflects the interaction of the query gene with the ~300 array genes also present in the chemical genetic interaction profiles.

#### Predicting a compound’s genetic targets

We predict the genetic targets (GTs) of a compound by calculating the similarity between the compound’s CG profile and the GI profile of each potential genetic target. However, common similarity measures such as Pearson and cosine correlation allow low-degree (low-intensity) CG profiles to correlate highly with GI profiles, despite their lack of signal. We therefore developed a target prediction score that gives preference to the genetic predictions made for high-degree over low-degree CG profiles.

This score, called the genetic target (GT) score, is calculated for each compound-GT pair as the dot product of the compound’s CG profile and the GT’s L_2_-normalized GI profile:

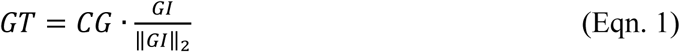

where *GT* is the genetic target score for the compound-GT pair, *CG* is the chemical genetic interaction profile of the compound, and *GI* is the genetic interaction profile of the genetic target. Using this score, higher degree (higher intensity) CG profiles will tend to have higher GT scores than will lower degree CG profiles, while the GI profile degree of each genetic target exerts no influence on its GT scores.

#### Predicting the biological process targets of compounds

We developed a method to predict the biological process targets (process targets, or PTs) of a compound, which combines the results of three different PT prediction methods. Each of these methods utilizes a similar scheme. First, GTs are mapped to PTs using propagated Gene Ontology biological process terms. Second, for each compound-PT pair, a z-score and empirical p-value are derived by calculating a statistic (mean or sum) on the GT scores for genes in the PT and comparing it to an appropriate null distribution. The specific details of each step are described in the following sections.

#### Defining the process targets

The process targets are a subset of terms from the “biological process” branch of the Gene Ontology annotations (http://geneontology.org/). The mapping of *S. cerevisiae* genes to biological process terms was obtained from the *Saccharomyces cerevisiae* Genome Database (http://www.yeastgenome.org/). Terms were propagated using “is_a” relationships, meaning that a gene annotated to term A was additionally annotated to term B if term A (the child term) had an “is_a” relationship with term B (the parent term). After propagation, terms with 4-200 genes annotations from the set of 1505 GTs included in our genetic interaction data were selected to become the “process targets.”

#### Process-based z-score and p-value

A z-score was computed to reflect the strength of a compound’s prediction to each candidate PT relative to control compounds’ predictions to the same PT. For a compound-PT pair, the relevant statistic used to calculate the z-score is the sum of the compound’s GT scores for genes in the PT (the “PT-sum”). The z-score for each compound-PT pair was calculated using equation 2, where *S* is the PT-sum for the compound-PT pair, *S¯* is the mean of the PT-sum values for that PT across all control CG profiles, and *σ* is the standard deviation of the PT-sum values for that PT across all control CG profiles.

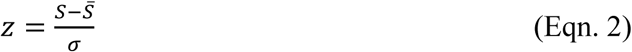

An empirical p-value was also derived to accompany this z-score. Specifically, for a given compound-PT pair, the empirical p-value was the fraction of times a control CG profile generated an equal or larger PT-sum than did the compound’s CG profile for that PT.

#### Compound-based z-score and p-value

A second z-score was computed to measure the specificity of each compound’s prediction to the candidate PT. The z-score for each compound-PT pair was calculated using equation 3, where s is the mean of the compound’s GT scores for genes in the PT, *N* is the number of genes annotated to the PT, *μ* is the mean of all GT scores for the compound, and *σ* is the standard deviation of all GT scores for the compound.

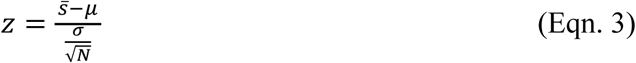

An empirical p-value was derived to accompany each z-score, providing for a significance-based comparison of the results with those from other z-score based metrics. To generate the null distribution for computing this p-value, the compound’s GT scores were randomly shuffled with respect to the GT labels 10,000 times. This preserves the mapping between GTs and PTs across the randomizations, and thus accounts for any effects due to the Gene Ontology structure. For a given compound-PT pair, the p-value is the fraction of times that the z-score calculated from the shuffled set of GT scores is equal to or larger than that calculated from the unshuffled scores.

#### Computing a combined z-score and p-value for compounds and their process targets

Three different sets of (z-score, empirical p-value) were computed for each compound-PT pair. One of these sets of values consisted of the compound-based z-score and p-value. Each of the other two sets was composed of a process-based z-score and p-value, with one z-score/p-value combination derived from 5724 experimental negative control profiles and the other z-score/p-value combination from 50,000 resampled CG profiles. The 5724 control profiles were generated by growing the yeast diagnostic mutant pool in the absence of a compound but in the presence of the compound’s solvent, DMSO (a negative control), and the 50,000 resampled profiles are described in more detail in the “Assessing the false discovery rate of process target predictions” section that follows.

To derive a single measure of significance, the largest (least significant) p-value of the three methods described above and its corresponding z-score were used as the final compound-PT score, as it represents the most conservative estimate of the strength and significance of a compound’s prediction to a PT.

### Assessing the false discovery rate of process target predictions

We assessed the false discovery rate of our process target predictions by predicting the process targets of control CG profiles and comparing the significance of those predictions with process target predictions made for the compound CG profiles. Two sets of control CG profiles were used: the DMSO condition CG profiles and the resampled CG profiles. 50,000 randomly resampled CG profiles were constructed by selecting one CG interaction score from each strain across all CG profiles in that screen. In this manner, the variance of each strain (especially, the tendency of certain strains to “drop out” of the pool) was captured in these resampled CG profiles.

For each set of compound and control conditions, the following procedure was employed to count the number of discoveries for FDR calculation: at every p-value, the number of conditions whose significant prediction was less than or equal to that p-value was counted; to enable a fair comparison between compound and control condition counts, the control condition counts were multiplied by a scale factor such that the total number of control conditions matched the total number of compound conditions.

A continuous false discovery rate estimate was then calculated as a function of the p-value for each type of control condition (DMSO and resampled CG profiles), by dividing the adjusted number of predictions made for the control condition by the number of predictions made for the compound condition at each p-value. Any resulting FDR values that were greater than 1 were adjusted to 1. The FDR derived from the resampled CG profiles was used for all subsequent analyses, as it was the more conservative (larger) of the two FDR estimates. From each screen, we identified a subset of high confidence predictions and combined them to generate the “high-confidence set” (HCS) (RIKEN: FDR ≤ 25%; NCI/NIH/GSK: FDR ≤ 27%).

To assess the performance of predictions, we identified known compounds with described modes-of-action present in our high-confidence prediction set (“gold standard compounds”). If the predicted process was functionally related to the known mode-of-action, we considered this a successful prediction (**Supplementary Table 20**).

### Characterizing the contribution of strains with high and low chemical-genetic interaction degree to process-level target prediction

We also characterized the contribution of the highest and lowest-degree strains to process-level predictions, this time by removing the 15% highest or lowest-degree strains before predicting process-level targets. The degree of a strain was defined as the number of interactions with CG score absolute value ≥ 2.5 it possessed across the RIKEN subset of compound-derived chemical-genetic interaction profiles (no DMSO or resampled profiles). After removing 40 of the highest or 41 of the lowest-degree strains (out of the 275 strains that overlapped with the *S. cerevisiae* genetic interaction network array strains, (**Supplementary Table 8**), process-level targets were predicted as described in “Predicting compounds’ modes of action based on chemical-genetic and genetic interaction profiles” and “Assessing the false discovery rate of process target predictions.” Comparisons regarding the number and identity of discovered compounds, and the identity of their predictions, were performed to determine the roles that high and low chemical-genetic interaction degree strains played in predicting process-level targets.

While the removal of low-degree strains had little effect on the identity of discovered compounds and their predictions, the removal of high-degree strains had noticeable effects. The “no-low-degree” profiles led to discovery of 927 bioprocess-level target predictions, 794 of which matched the original RIKEN “all-strain” predictions (94% of the 848 original RIKEN high confidence set, or HCS) (**Supplementary Table 8.1**). In contrast, the “no-high-degree” profiles led to the discovery of only 667 high confidence bioprocess-level target predictions, most of which overlapped with the RIKEN HCS (537 compounds, or 63% of the RIKEN HCS). In addition, the predictions derived from “no-low-degree” profiles tended to match the predictions of in the RIKEN HCS (602/794, or 76%, of “all-/no-low-degree” compounds shared predictions with Jaccard ≥ 0.25), while the predictions derived from “no-high-degree” profiles were less consistent (168/667, or 31%, of “all/no-high-degree” compounds shared predictions with Jaccard ≥ 0.25).

The importance of high-degree strains to bioprocess-level predictions was further confirmed by examining the identities of the predicted processes. While removing high-degree strains does not destroy the performance of bioprocess-level predictions, it does substantially change the distribution of the most frequently-predicted bioprocesses and reduce prediction accuracy for some well-characterized compounds. After removing high-degree strains, the top predicted bioprocess by far was “spindle assembly,” followed by other microtubule and cell cycle-related processes, and finally, bioprocesses related to localization, pH and ATP, glycosylation, and DNA damage/repair **(Supplementary Table 8.2)**. For three well-characterized compounds, the removal of high-degree strains substantially reduced prediction specificity for tunicamycin, altered predictions of rank 3 and below for benomyl, and left the predictions for MMS essentially unchanged (**Supplementary Table 8.3**). In contrast, removing low-degree strains had little effect on either the distribution of process-level predictions in the high-confidence set or the highest-confidence predictions for benomyl, MMS, and tunicamycin.

### Characterizing the respective contribution of negative and positive interactions to process-level target prediction

Using the RIKEN NPDepo high-confidence set of compounds, we characterized the contribution of positive and negative chemical-genetic interactions to our process-level predictions. First, chemical-genetic interaction profiles containing either only positive or only negative interaction scores were generated. Process-level targets were then predicted using these “positive-only” or “negative-only” profiles as described in “Predicting compounds’ modes of action based on chemical-genetic and genetic interaction profiles” and “Assessing the false discovery rate of process target predictions.” We then compared the number and identity of the compounds discovered, and the identity of their predictions, between “positive-only,” “negative-only,” and “all-interaction” prediction sets to determine which side(s) of the chemical-genetics interaction profiles were important for predicting perturbed processes.

Two schemes were employed to generate the “positive-only” and “negative-only” chemical-genetic interaction profiles and their subsequent process-level predictions. Scheme 1 profiles showed how all negative and all positive interaction scores contribute to process-level predictions, and scheme 2 profiles accounted for biases that could have occurred due to differences in the number of positive vs. negative interactions in the scheme 1 profiles. To generate “negative-only” profiles under scheme 1, the positive scores in all compounds, DMSO control profiles, and resampled profiles were set to zero; conversely, “positive-only” profiles under this scheme were generated by setting all negative scores to zero. To generate the “positive-only” and “negative-only” profiles under scheme 2, an equal number of scores with absolute value ≥ 1 were selected from the extreme positive or negative ends, respectively, for each compound, DMSO, and resampled profile.

“Negative-only” and “positive-only” chemical-genetic interaction profiles led to the identification of a substantially different sets of “high-confidence” compounds (at least one prediction with FDR ≤ 25%), with the “negative-only” profiles reproducing the “all-interactions” high confidence set much better than did the “positive-only” profiles. Both high confidence sets derived from “negative-only” profiles from scheme 1 (all scores) and scheme 2 (equal number of positive vs. negative scores) possessed roughly the same number and identity of compounds when compared to the “all-interaction” high confidence set (**Supplementary Table 9)**. Specifically, 85% (723/848) and 81% (689/848) of the high confidence compounds identified using all interactions were discovered using scheme 1 “negative-only” profiles (“negative-all/all” comparison) and scheme 2 “negative-only” profiles (“negative-equal/all” comparison), respectively. While the high confidence set derived from scheme 1 “positive-only” profiles was similar in size to the “all-interactions” high confidence set, the compounds in both scheme 1 and scheme 2 “positive-only” high confidence sets had much lower overlap with the “all-interactions” high confidence set (345/848, or 41% – “positive-all/all” comparison, and 183/848, or 22% – “positive-equal/all” comparison, respectively).

In addition to driving the discovery of the same compounds that were in the “all-interactions” high confidence set, negative chemical-genetic interactions also drove the discovery of the same predictions for these compounds. For example, 68% (494/723) of the “negative-all/all” co-identified compounds and 47% (326/689) the “negative-equal/all” co-identified compounds had a Jaccard coefficient of ≥ 0.25 for their predictions. In contrast, only 17% (58/345) of the “all/positive-all” and 3% (6/183) of the “all/positive-equal” co-identified compounds met this criterion for the similarity of their predictions, suggesting that even for compounds where predictions were made, the predicted modes of action were largely different. From this evidence, negative chemical-genetic interactions are clearly the primary driver of genetic interaction-based target predictions.

In addition, two lines of evidence suggest that the predictions made using only positive chemical-genetic interactions are of lower quality than those derived from all or only negative interactions. First, we observed that the predictions from positive chemical-genetic interactions were overwhelmingly biased toward GO terms related to RNA splicing/processing and cell cycle/mitosis, while those from all or only negative interactions were more diverse (GO terms related to cellular localization, chromatin organization and transcription, cell wall, vesicle-mediated transport, pH regulation, protein degradation, microtubules and cytoskeleton, etc., in addition to cell cycle/mitosis) (**Supplementary Table 9.1**). Second, we observed that in the set of predictions derived from only positive interactions, three well-characterized compounds (benomyl, MMS, tunicamycin), whose known mechanisms of action are well-captured by process-level predictions based on either all or only negative interactions, both 1) failed to make the high confidence compound list and 2) did not show predictions consistent with known mechanisms (**Supplementary Table 9.2**).

### Visualizing the relationship between compound bioactivity and inclusion into the high confidence set

We assessed the fraction of compounds in the high confidence set as a function of bioactivity, which can also be thought of as the probability that a compound will be in the high-confidence set given its bioactivity. The bioactivity (percent growth compared to DMSO) and high confidence set status (true/false, respectively set to 1/0 for analysis) for each compound were extracted from **Supplementary Table 4**. A loess curve was then fit through the 1/0 high-confidence status values with respect to the bioactivity values, using a span of 0.1 and least-squares fitting with a polynomial degree of 2. The curve on the plot was drawn at points 2.5 units apart, starting at the smallest observed bioactivity value (**Supplementary Fig. 6**).

### Determining functional distributions of compound collections

#### Generating the background set of chemical genomic profiles

To account for biases in the distribution of process predictions introduced by our discovery pipeline, we generated a set of “background” chemical genomic profiles. Each background profile was a high-signal GI profile with noise added based on the variance of each strain across all GI profiles (Gaussian, *μ* = 0, *σ* = 2 × *σ_strain_*). Each of these 4515 profiles (3 for each of 1505 GI profiles) simulated a compound that targets one gene. This enabled the estimation of any functional biases introduced by our GI-based discovery pipeline.

#### Computing distributions of process predictions for each compound class

We calculated the proportion of each compound class that was predicted to each process term. (**Supplementary Table 21**). Those proportions were then compared to the proportion of the background profiles predicted to each process using a proportion test in R (**Supplementary Table 22**). To sort from the most significant enrichment to the most significant depletion compared to the background, p-values from the proportion test were modified such that p-values from proportions greater than the background ranged from 0 to 1, and p-values from proportions smaller than the background ranged from 2 to 1. Using a ranksum analysis with the modified p-values as the input, we determined, for each class, if processes that mapped to each functional neighborhood were predicted more or less frequently than in the background set. Rank-sum p-values were Bonferroni-corrected and visualized as a heatmap (**Fig. 4b**).

### Compound diversity sets for functional neighborhoods

We assigned all the compounds associated with a specific functional neighborhood to a single cluster and split up the cluster recursively to form clusters of more similar compounds. At any recursive step, we determined the cluster with the lowest average within-cluster chemical genomic similarity and divided the cluster into two new clusters using K-means clustering. We stopped generating new clusters right before our algorithm would generate at least two individual clusters exceeding our predefined limit for the maximum average between-cluster chemical genomic similarity (cosine similarity of 0.3). We repeated the algorithm 1000 times for each neighborhood and selected, from each cluster, the compound with the strongest prediction as a candidate for our diversity set. We finally sorted all our candidates across all the repetitions from the most frequent to the least frequently occurring. To define the compound diversity set, we selected from this ranked list as many top candidates as were needed to cover all the clusters in at least 50% of the repetitions.

### Comparison with other chemical genomic datasets

An independent set of whole-genome chemical genomic screens have been performed previously by Lee et al., (2014) and Hoepfner et al., (2014)^14,15^. These studies interrogated 3,239 and 2,923 compounds, respectively, and they were performed using both a heterozygous and homozygous diploid deletion mutant profiling platform. The homozygous diploid deletion mutant profiling platform is comparable to the chemical-genetic analysis we carried out with haploid deletion mutants. Our study shares 145 compounds in common with the Lee et al. study and 31 compounds in common with the Hoepfner et al. study. In particular, all three studies reported an overlap of 9 compounds.

Comparisons were made between our chemical-genetic interaction scores with the Hoepfner et al. median absolute deviation logarithmic scores, and with the Lee et al. fitness defect scores (multiplied by −1), such that the chemical-genetic interaction profiles were restricted to the 277 genes common between the three studies. For the nine shared compounds (**Supplementary Table 6**), our study shows an average Pearson correlation coefficient (PCC) of 0.29 with Lee et al., and 0.38 with Hoepfner et al. whereas Lee et al. and Hoepfner et al. show a PCC of 0.22. Thus, our study shows significant agreement with both the Lee et al., study (p-value: 5 × 10^-7^) and the Hoepfner et al. study (p-value: < 1 × 10^-8^).

We also compared the members of the compound diversity sets derived from our RIKEN and Clinical screens to the major chemical-genetic signatures defined in Lee et al. and found favorable overlap of the chemical space occupied by compounds from both studies. After computing PCC between each diversity set compound and each compound from Lee et al. that was annotated to a major signature, we observed that all 45 major Lee et al. signatures contained at least one compound that was significantly similar to a compound in both diversity sets (PCC > 0.2, one-sided test, p-values obtained by shuffling the profile gene labels 10,000 times followed by Benjamini-Hochberg correction, FDR < 0.05) and that most of the compounds in the RIKEN and Clinical diversity sets contributed to this overlap (123/130 unique RIKEN and 187/214 Clinical compounds) (**Supplementary Fig. 15a**). When applying a more stringent PCC threshold (PCC > 0.4), only 18 and 12 (out of 45) Lee et al. major signatures are covered by 32 and 39 compounds from the RIKEN and Clinical diversity sets, respectively.

In addition, we mapped the Lee et al. major signatures to our bioprocesses and found that many of these mappings agree functionally (**Supplementary Fig. 15b**). After computing PCC between the profiles of each high confidence compound and each compound from Lee et al. that was annotated to a major signature, we annotated each correlation > 0.3 to a major signature/bioprocess pair (the bioprocess annotation for each high confidence compound was based on its best process prediction). For each major signature/bioprocess pair, we then counted the number of unique Lee et al. and high confidence compounds, respectively, that contributed to these correlations. We normalized these counts by the size of their respective major signature or bioprocess and multiplied the resulting fractions together to derive a confidence score that deemphasizes major signature/bioprocess pairs for which a very small number of compounds annotated to the major signature (or bioprocess) is responsible for most of the correlations to the compounds in the bioprocess (or major signature). A table that maps each Lee et al. major signature to its most confident bioprocess is provided (**Supplementary Table 23**), as is a table that maps each Lee et al. signature to any bioprocess with which it shared at least one profile correlation > 0.3 (**Supplementary Table 23.1**). Both tables are sorted by confidence in descending order. Agreement between the Lee et al. major signatures and our bioprocess annotations was encouraging; specifically, Golgi (Lee et al.) mapped to Vesicle traffic (this study), ubiquinone biosynthesis & proteasome (Lee et al.) to Protein Degradation (this study), ergosterol depletion effects on membrane (Lee et al.) to Metabolism & Fatty Acid Biosynthesis (this study), and DNA damage response (Lee et al.) to DNA Replication & Repair (this study). Overall 43/45 major chemical-genetic signatures possessed at least one compound with PCC > 0.3 to a compound in our study and therefore could be mapped to a bioprocess; however, mappings derived from a very small number of compounds in either member of the pair should be interpreted with more caution.

### Identifying structural motifs contributing to functional enrichments

To identify structural motifs that drove specific functional neighborhood enrichments, we performed discriminative molecular substructure mining on the RIKEN HCS set of compounds using the MoSS tool^67^. Using the proportion of each compound class that was predicted to each process term (see “Computing distributions of process predictions for each compound class”), we selected only process terms that had a significantly higher proportion of predictions in at least one compound class versus the GI background (proportion test in R, Bonferroni-corrected). Then, for each process term, we identified substructures that occurred at least twice as frequently in compounds with high confidence predictions to that process term (the “active” set) versus compounds that did not have high confidence predictions to that term (the “inactive” set). This discriminative mining was performed twice per process term: once by drawing the inactive set of compounds from all screened compounds in the RIKEN NPDepo, and once by drawing the inactive set from all NPDepo compounds in the HCS. By selecting the minimum of these two enrichments, we sought to control for bias in the distribution of substructures in the inactive compounds. The information about the substructures and their enrichments was compiled across all experiments. The final output is a table of substructures that show enrichment for a particular functional category (**Supplementary Table 14**).

### Localization enrichments

We sought to determine if the compounds in particular collections exhibited bias in the localization of their targets. Using the proportion of each compound class that was predicted to each process term (see “Computing distributions of process predictions for each compound class”), we selected process terms that had significantly higher (enriched) and lower (depleted) proportions of predictions versus the GI background (proportion test in R, Bonferroni-corrected). For each compound collection, two gene lists were assembled, each representing the union of the genes annotated to either enriched or depleted (*p*_bonf_ ≤ 0.05) process terms.

A hypergeometric test was performed to determine which of these gene lists were enriched for genes annotated to specific cellular components. P-values were Bonferroni-corrected. Gene annotations to cellular compartments were obtained from Huh et al. 2003^68^, Koh et al. 2015^69^, and the yeast GO slim cellular compartment annotations (http://www.yeastgenome.org/). The background set of genes for all hypergeometric tests was the set of 1499 query genes with GO process annotations from the high-degree genetic interaction dataset.

### Flow cytometry based global validations of targeted processes

67 compounds with process target predictions mapping to G1-phase arrest, S-phase arrest, or G2-phase arrest flow cytometry phenotypes (based on Yu et al. 2006^40^) were selected from the high confidence set. Compounds that ultimately mapped to multiple cell cycle phenotypes via their process target predictions were removed from consideration. For each cell cycle phenotype, the 50 compounds with the highest overlap of 1) gene targets driving the process prediction that mapped to the phenotype and 2) the genes directly annotated to the phenotype (Yu et al., 2006) were selected. Compounds were then manually selected from these lists based on their bioactivity, as compounds with higher bioactivity were assumed more likely to induce a cell cycle phenotype.

Cultures of the control strain (y13206) were grown to early log phase (0.4 OD) in YPGal (1% yeast extract, 2% peptone, 2% galactose). 250 μL per well of the starting culture was aliquoted into a 96-well block. The cultures were treated with 10 μg/mL of each compound and incubated at 30 °C for 2-3 h. We included the compounds hydroxyurea, MMS, nocodazole, and tunicamycin as controls known to arrest cell cycle in G1, S, G2 and post-G2 respectively. From each culture, 200 μl was transferred into a new 96 well plate, pelleted at 2000 rpm for 5 min. Pellets were resuspended in 20 μL of 50 mM Tris-Cl (pH 8.0), 50 mM EDTA buffer. 160 μl of cold 99% EtOH was added to the wells. Cells were pelleted at 4000 rpm for 2 min at RT, resuspended in RNAse A solution (50 mM Tris-Cl pH 8.0, 0.4 mg/mL RNAseA), and incubated for 2 h at 37 °C. Cells were pelleted at 4000 rpm for 2 min at RT, and 50 μL of proteinase K solution was added (50 mM Tris-Cl pH 7.2, 200 mM NaCl, 78 mM MgCh, filter sterilized). The cells were then incubated for 50-60 min at 50 °C. Cells were pelleted at 4000 rpm for 2 min at RT, and resuspended in 55 μL of FACS buffer (200 mM Tris-Cl pH 7.5, 200 mM NaCl, 78 mM MgCh, filter sterilized). In a new 96 well plate, 180 μL of SYBR Green solution (2X SYBR Green, 50 mM TrisCl pH 7.2) was added to each well. 20 μL of fixed cells from the previous step was added. The plate was then processed via high-throughput flow cytometry as described in Yu et al., 2006. The voltage of the green channel was adjusted so that on the linear scale the 1C peak and the 2C peak were well spaced, the 1C peak was away from the vertical axis. The FSC-A vs FL1-A was used to gate out aggregates and dead cells. The final histograms have FL1-A on the x-axis (area of the green channel).

Cell cycle phenotypes were called by drawing thresholds based on 46 control DMSO profiles, on either the percent of cells in S phase (%S) or the ratio between the percentages of cells in G1 (1C peak) vs. G2 (2C peak) phase (G1/G2 ratio). Specifically, the mean and standard deviation were computed for both the %S and the G1/G2 ratio in the DMSO control samples. These values were used to convert the corresponding values from the treatment compounds into z-scores. A phenotype was called if the z-scores of both replicates passed the appropriate z-score threshold of either 1.5 or −1.5. The specific thresholds for phenotypes calls were as follows: a 1C phenotype was called if G1/G2 ratio > 1.196; a 2C phenotype was called if G1/G2 ratio < 0.809; and an S phenotype was called if %S > 19.5%.

Enrichments and p-values were computed empirically by shuffling the phenotypes associated with the compounds and counting the number of cell cycle phenotypes associated with each prediction in the shuffled data (100,000 randomizations). Compound identities were preserved during the randomization, such that both replicates of a compound were associated with the same cell cycle phenotype prediction after each randomization. Enrichments were computed by dividing the number of calls observed from the real data by the average expected number of calls for each combination of predicted and observed phenotype (averaged over all compound-predicted phenotype randomizations). In a similar fashion, empirical p-values were computed for each combination of predicted and observed phenotype by counting the fraction of randomizations that produced the same or larger number of calls.

### Multi-parameter validation of cell wall targeting compounds

For the adenylate kinase (AK) cell leakage assay, an overnight culture of the drug hypersensitive yeast strain (y13206) in log phase was harvested and washed twice with fresh YPGal medium. The final pellet was resuspended in 1 mL fresh YPGal. Fifty microliters of cell suspension (~1×10^6^ cells), 1% DMSO, 10 μg/mL of each test compound was added in individual wells of 96-well culture plate containing YPGal medium to a final volume of 100 mL, mixed by pipetting and incubated at 25 °C for 4 h (n=3). The plate was equilibrated to room temperature for 30 min and the contents were transferred into a luminescence compatible 96-well white-walled plate. Next, 100 μL of ToxiLight AK reagent (Lonza) was added to each well and incubated at room temperature for 30 min, and luminescence was measured with a Wallace ARVO SX 1420 Multilabel Counter (Perkin Elmer Life Sciences). Hit compounds resulted in more than 20000 units. Cells were stained with the glucan stain aniline blue and the chitin stain calcofluor white as described previously^41^, and hits assessed by irregular glucan or chitin staining detected by eye. Treated cells were analyzed by high-dimensional morphometric analysis (CalMorph) as described previously (n=5)^70^. A neck width and morphological noise (heterogeneity) was determined as described previously^41^.

### Zymolyase sensitivity assay

Zymolyase sensitivities were tested as described previously^71^ with slight modifications. Yeast cells (y13206) were grown in YPGal until log phase (~4×10^7^ cells/mL), and 50 mL of aliquot was transferred into fresh 150 mL YPGal containing test compounds in 96-well microtiter plate (10 or 40 mg/mL for test compounds, as for controls: 2.5 mg/ml for echinocandin B, 30 mM for hydroxyurea, 1% for DMSO). The cell-containing plate was incubated at 25 °C for 4 h with shaking. After incubation, cells were washed twice with 10 mM Tris-HCl (pH7.5), and resuspended to zymolyase solution (0.94 mg/mL of Zymolyase 100T (Seikagaku) in 10 mM Tris-HCl (pH 7.5)). Cell suspensions were incubated at 30 °C, and OD_600_ values were measured for 1 h after the addition of zymolyase with plate reader (SPECTRAmax plus384, Molecular devices). In each sample, OD_600_ values were standardized at time 0 to equal 1 (or 100%).

### Cell cycle analysis of NPD5925

Y13206 cells were grown to mid-log phase in YPD, and a sample of this asynchronous population was saved for later analysis. The cells were treated with alpha factor and incubated for 2.5 hours at 30 °C, and a sample of the alpha factor-arrested population was saved for later analysis. Pronase and test compounds were added to the remaining arrested population. We tested DMSO (2%), hydroxyurea (0.2 M), MMS (0.03%), and NPD5925 (20 μg/mL). The treated cells were incubated for 1 h and then prepared and analyzed via flow cytometry as described above.

### Tubulin inhibition assay and assessing predictive power

We carried out *in vitro* tubulin polymerization assays using the cytoskeleton fluorescent based porcine tubulin polymerization assay (Cytoskeleton Inc, USA) following manufacturer specifications. We used 10 μg/mL of test compound for each assay. We tested the control compounds nocodazole, paclitaxel, and the predicted tubulin targeted compound NPD2784 versus a DMSO solvent control.

### Identifying compounds with multiple, unique mechanisms of action

We devised an algorithm to prioritize compounds from the RIKEN HCS whose chemical genetic (CG) interaction profiles appeared to be a combination of multiple, diverse genetic interaction (GI) profiles, indicating that they exert their effects via multiple, unique mechanisms of action. For a compound, we first constructed profiles reflecting the mean contribution of each strain in its CG profile to each of its process target (PT) predictions. Then, the initial cluster of “mean contribution profiles” was seeded with the profile from the highest confidence PT prediction. To complete the clustering, the mean contribution profiles from progressively lower-confidence PT predictions were either added to an existing cluster (if they possessed a Pearson correlation coefficient of ≥ 0.5 with a profile in that cluster) or used to seed a new cluster. Compounds were prioritized if they possessed two clusters of mean contribution profiles with very low average similarity between them, suggesting that two distinct signals in the GI network contributed to the signal observed in their CG profiles.A set of contribution profiles was generated for a compound and one of its PT predictions by taking the element-wise product of the compound’s CG profile and the L_2_-normalized GI profile of each gene that drove the PT prediction (genes with genetic target score ≥ 2 and were annotated to the PT, which are shown in columns “driver_common” and “driver_score” in **Supplementary Table 7**. The “mean contribution profile” for one compound and PT prediction was calculated as the strain-wise mean across all of the contribution profiles associated with that compound and one PT prediction. GI profiles were from the set of high-signal genetic interaction profiles.

### Staining of cells with NPD5925

Log phase yeast cells (y13206) were fixed with 3.7% formaldehyde solution. The fixed-cell suspension was centrifuged to make cells a pellet, and the pellet was mixed with the same volume of NPD5925 (1 mg/mL) and incubated at 25 °C for 30 min. Cells were washed twice with phosphate-buffered saline (PBS), and a small cell aliquot was mixed with mounting solution (90% glycerol, 9.975% PBS, 0.025% 0.1 N NaOH) containing *p*-phenylenediamine (1 mg/mL) and 4',6- diamidino-2-phenylindole (DAPI, 0.7 mg/mL). A prepared specimen was observed by fluorescent microscope (Axioimager M1, Carl Zeiss) with regular rhodamine or DAPI filter sets (Carl Zeiss). An intensity profile was extracted from cell images by ImageJ (http://imagej.nih.gov/ij/).

### Adenylate kinase (AK) assay of NPD5925

An overnight culture of yeast strain (y13206) in log phase was harvested and washed twice with fresh YPGal medium and the final pellet was resuspended in 1 mL fresh YPGal. Fifty microliters of cell suspension (~1×10^6^ cells), 1% DMSO, 30 mM hydroxyurea, 20 mg/ml Echinocandin B, and 40 mg/mL test compounds were added in individual wells of 96-well culture plate containing YPGal medium to a final volume of 100 ml, mixed by pipetting and incubated at 25 °C for 4 h. The plate was equilibrated to room temperature for 30 min and the contents were transferred into a luminescence compatible 96-well white-walled plate. Next, 100 μL of ToxiLight AK reagent (Lonza) was added to each well and incubated at room temperature for 30 min, and luminescence was measured with a Wallace ARVO SX 1420 Multilabel Counter (Perkin Elmer Life Sciences).

### Assessing potential targets of NPD5925 for pleiotropy between DNA and cell wall processes

The chemical genetic interaction profile was compared against high confidence genetic interaction profiles using a genetic interaction normalized cosine score (genetic target score, Eqn. 1 above). The top ten high confidence genes were displayed alongside the chemical genetic interaction profile.

For each of the high confidence GO process predictions for NPD5925, the genetic interaction profile of the drivers of that GO process prediction, high confidence genes with a genetic target score above 2 annotated to the enriched GO process, were combined using the following to form a GO process specific importance profile,

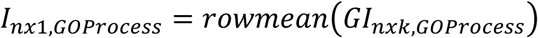

where k is the number of genes driving the GO process prediction. A GO process driven chemical genetic interaction profile is then derived with:

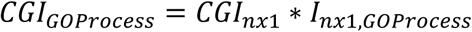

The GO process driven chemical genetic interaction profile is then compared against genetic interaction profiles in high confidence using the genetic target score, and the top ten high confidence genes were displayed along the GO process driven chemical genetic interaction profile.

